# Mimicking Human EED Variants in Drosophila: A Promising Strategy to Analyse Human EED Variant Function

**DOI:** 10.1101/2024.02.18.580839

**Authors:** Sharri S. Cyrus, Sònia Medina Giró, Tianshun Lian, Douglas W. Allan, William T. Gibson

**Author notes:** Douglas W. Allan is the corresponding author regarding *Drosophila* experiments and strains and William T. Gibson is the clinical corresponding author.

## Abstract

The Polycomb Repressive Complex 2 is an epigenetic reader/writer that methylates histone H3K27. Rare germline partial loss of function (pLoF) variants in core members of the complex (EZH2, EED, SUZ12) cause overgrowth and intellectual disability syndromes, whereas somatic variants are implicated in cancer. However, up to 1% of the general population will have a rare variant in one of these genes, most of which would be classified as variants of uncertain significance (VoUS). Towards screening these VoUS for partial LoF alleles that may contribute to disease, here we report functional assays in *Drosophila* to interrogate *EED* missense variants. We mimicked the amino acid change(s) of *EED* variants into its *Drosophila* ortholog, *esc*, and tested their function. Known likely benign variants functioned wildtype and known pathogenic variants were LoF. We further demonstrate the utility of this calibrated assay as a scalable approach to assist clinical interpretation of human *EED* VoUS.

## Introduction

It is estimated that at least 65% of the human clinome (i.e. disease-causing genes) have a functional ortholog in *Drosophila*.^1^ Despite gross morphological differences between fruit flies and humans, the molecular mechanisms that control fundamental biological processes, physiology and cellular development are conserved between the two species.^1^ This offers opportunities to unravel mechanisms for human disease, and also to derive insights into the functional effects of individual variants. *Drosophila* as an organism has been extensively studied for more than a century,^2^ and thousands of genes have documented mutant phenotypes that can be used as readouts when studying the functional impact of naturally-occurring variants in conserved human proteins. Such assays are easily scalable owing to *Drosophila*’s fecundity, short generation time and cost-effective genetic tractability. This scalability allows for large numbers of variants to be screened, which is ideal to interrogate the many variants of uncertain significance (VoUS) seen in human populations. Here, we describe assays that can be used to interrogate human *EED* coding variants with clinical predictive value.

*EED* is a member of the well conserved Polycomb Repressive Complex 2 (PRC2). Germline heterozygous partial loss of function (pLoF) mutations cause the rare Cohen-Gibson syndrome (COGIS), whereas somatic LoF variants have been implicated in haematological malignancies.^3,4^ To date, only a few variants that cause COGIS have been assessed functionally. For example, the R302S and R236T mutations seen in two individuals with COGIS resulted in a reduction of H3K27me3 levels in human 293 T-REx cell lines.^3^ A similar result was observed for R236T in a patient-derived lymphoblastoid cell line.^3^ However, most COGIS variants remain unexamined, and no single model has been established to compare the function of multiple EED variants. Additionally, the widespread use of sequencing technologies has made genomicists aware of the ubiquity of VoUS in many genes in every individual’s genome. This underscores the growing need for functional assays to resolve these variants, especially for disease-associated genes. As diagnostic exome sequencing becomes more cost-effective,^5^ and as rare variant burden testing eclipses genome-wide SNP arrays for disease associations in human populations,^6^ reclassification of VoUS into the likely benign (LB) and likely pathogenic (LP) categories will become increasingly important for accurate interpretation of exome-level and genome-level data. Because rare and ultra-rare VoUS may never be re-classifiable on the basis of allele frequencies and/or computational predictions, functional data are needed to fill the classification chasm.

Here, we mimic *EED* human variants in the fly’s ortholog, *esc*. We then assess the ability of these variants to functionally replace *esc*, using well-characterized phenotypes as readouts. Numerous studies have documented the utility of such a mimetic approach to demonstrate a human variant’s functional impact and inform disease mechanism.^7–10^ Indeed, recent work showed the utility of such a mimetic approach by identifying a gain of function impact for a rare human EZH1 variant in the context of an identical amino acid change in the orthologous *E(z)* fly gene a fly model.^11^ Here, we demonstrate that known pathogenic (P) and LB EED variants faithfully recapitulate their relative function when mimicked in the *esc* gene, allowing for successful calibration of an assay to test *EED* VoUS. This method appears useful to test variants in the context of established mutant phenotypes for the *Drosophila esc* gene, and to avoid the challenge of determining parameters for human EED function in *Drosophila* before an assay can be employed.

Mutations in *Drosophila* Polycomb group genes cause homeotic phenotypes, due to the misexpression of homeotic (Hox) genes.^12–14^ Misexpression of these genes results in homeotic transformations of segments and structures.^15^ Mutations in *esc* can cause developmental lethality or an extra sex comb phenotype in adult males, wherein sex combs are found ectopically on the 2^nd^ and/or 3^rd^ pair of legs in males.^12,16^ These phenotypes are well-established, are easily-screened, and can be used to demonstrate if *esc* variants that mimic human *EED* orthologous variants (referred to as *esc* mimetics) confer functional impairments on the Esc protein.

In one approach, we exploited an existing CRIMIC *T2A::GAL4* insertion within a coding intron of the *esc* gene,^17^ which replaces Esc protein expression with GAL4. This served as a platform for both ϕC31-integrase Recombinase-Mediated Cassette Exchange (RMCE) of the *T2A::GAL4* cassette into multiple germline-integrated *T2A::esc* mimetic variant alleles (Figure 1), and also for GAL4-mediated expression of *UASz-esc* mimetic variants that we germline-integrated into the *attP2* site by ϕC31 integrase. In an alternate approach, we used homology-directed repair (HDR) CRISPR/Cas9 genome editing to make a heritable coding change in the native *esc* gene to mimic a specific EED variant. Using these approaches, we selected 22 *EED* variants for mimicry (Table 1/Figure 1): a mixture of LB and P/LP variants, VoUS, somatic LoF variants and a single null variant . For simplicity, all mimetic variants will be referred to by the EED amino acid position (not Esc position) throughout this report. Table 1 shows the amino acid change in fly Esc that mimics each human EED variant. We show that our assays can accurately discern LB *esc* mimetic variants from P/LP variants, satisfying key criteria of the ACMG recommendations for clinical interpretation of variant function in experimental assays.^18^ Having successfully calibrated this assay, we then tested the function of numerous EED VoUS and reclassified them as LB or LP. We believe that this approach can reclassify EED VoUS based on their relative function in a *Drosophila* model.

**Figure 1:**
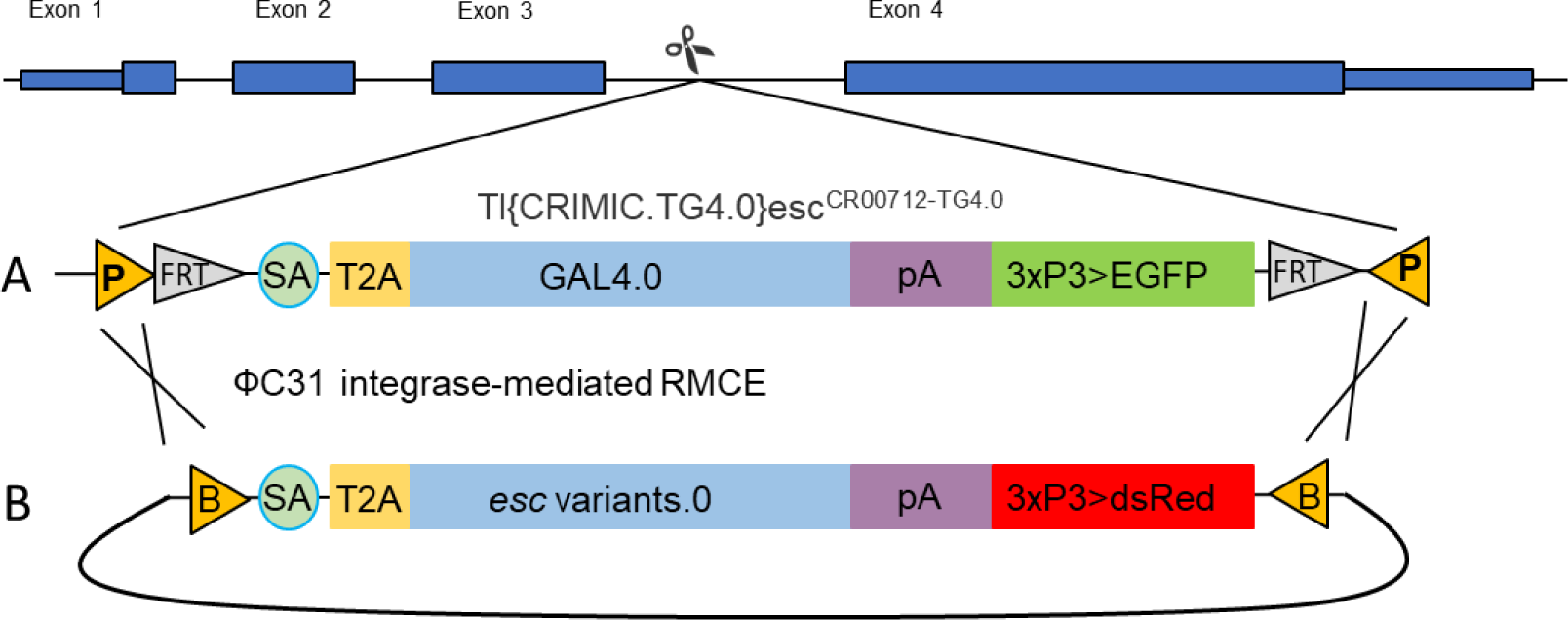
Generation of *esc* mimetic variants by Recombinase Mediated Cassette Exchange. **A)** A swappable integration cassette (SIC) was inserted as an artificial exon (containing *cis*-inverted *attP* sites (P), flippase recombination target (*FRT*) sites, a splice acceptor (SA), a *T2A*::*GAL4* coding sequence in frame with 5’ *esc* coding sequence, an SV40pA terminator sequence (*pA*) and a green eye fluorescent marker (3xP3>EGFP)), into the intron by Cas9-mediated homology directed repair (HDR). The resulting CRIMIC allele generates a transcript that splices into the artificial exon and terminates transcription without splicing into the native exon 4. The resulting translated product encodes the N-terminus of the native fly protein, followed by an in-frame self-cleaving T2A peptide that releases an in-frame, full length GAL4 protein. This fly line (*esc::T2A::GAL4*) was generated by the Gene Disruption Project,^17^ and made available at the Bloomington *Drosophila* Stock Centre. **B)** We used the SIC as a host landing site to integrate sequences between *cis*-inverted *attB* sites (B) (within embryo-injected plasmids) into the *cis*-inverted *attP* sites, by ϕC31-integrase-mediated Recombinase-Mediated Cassette Exchange (RMCE). This is a high-efficiency, high-fidelity method to introduce *esc* mimetic gene variants (*esc* cDNA mutant) into the locus that would be expressed in the same spatiotemporal pattern as endogenous *esc*. Once the construct is integrated, T2A self-cleavage releases full-length mutant esc protein (instead of GAL4) driven by the endogenous *esc* promoter, and dsRed replaces EGFP as the selectable marker.

**Table 1:** Human EED Variants Tested. P:Published variant; ‘*’: ClinVar: criteria provided single submitter; **‘NP’**: ClinVar: no assertion criteria provided; **N/A**: Not assessed, absent in ClinVar. Allele counts in this Table are from gnomAD version 2.1.1, prior to the November 1, 2023 update to version 4.0.

| EED Variant | esc Orthologous Variant | Clinical Significance | ClinVar Annotation | References |
| --- | --- | --- | --- | --- |
| Likely Benign Variants |  |  |  |  |
| T50P | T32P | gnomAD: Allele count 81 | Likely benign* | 19 |
| I274V | I254V | gnomAD: Allele count 46 | Likely benign* | 19 |
| Likely Pathogenic/ Pathogenic Cohen-Gibson syndrome (COGIS) Variants |  |  |  |  |
| N194S | N174S | COGIS <sup>P</sup> | Pathogenic* | 20–23 |
| R236G | R216G | COGIS <sup>P</sup> | N/A | 20 |
| R236T | R216T | COGIS <sup>P</sup> (LoF) | Pathogenic<sup>NP</sup> | 3,23 |
| D237G | D217G | COGIS <sup>P</sup> | N/A | 20 |
| H258Y | H238Y | COGIS <sup>P</sup> (LoF) | Pathogenic<sup>NP</sup> | 21,23–25 |
| R302G | R282G | COGIS <sup>P</sup> Myeloproliferative neoplasms (LoF) | Pathogenic<sup>NP</sup> | 23,25,26 |
| R302S | R282S | COGIS <sup>P</sup> (LoF) | Pathogenic<sup>NP</sup> | 23,25,27 |
| 306R_307NdelinsTD | 286R_287NdelinsTD | COGIS <sup>P</sup> | N/A | 28 |
| A378V | A358V | COGIS | Likely Pathogenic* | 23 |
| LoF Variants in Haematological Malignancies |  |  |  |  |
| L240Q | L220Q | Refractory cytopenia with unilineage dysplasia (LoF) <sup>P</sup> | N/A | 4 |
| G255D | G235D | Atypical chronic myeloid leukemia (LoF) <sup>P</sup> | N/A | 29 |
| S259F | S239F | Acute lymphoblastic leukaemia (LoF) <sup>P</sup> | N/A | 25,30 |
| I363M | I343M | Myelodysplastic syndromes, CMML1 (LoF) <sup>P</sup> | N/A | 4,31 |
| Variants of Uncertain Significance |  |  |  |  |
| R52C | R34C | Hereditary breast ovarian cancer syndrome | Uncertain Significance* | 19,23 |
| T158M | T138M | gnomAD Allele Count:3 | N/A | 19 |
| F198L | F178L | Patient Derived | N/A |  |
| H258L | H238L | COGIS <sup>P</sup> | N/A | 21 |
| 306R_307NdelinsTD Component Variants |  |  |  |  |
| R306T | R286T | Pathogenic when in cis with N307D | N/A |  |
| N307D | N287D | Pathogenic when in cis with R306T | N/A |  |
| Null |  |  |  |  |
| L196P | L176P | null (LoF) <sup>P</sup> | N/A | 32,33 |

## Results

### A CRIMIC-based gene replacement strategy efficiently interrogates the function of *EED* variants

We used the *esc::T2A::GAL4* CRIMIC allele, *esc^CR^*^00712^*^-TG^*^4.0^, as a platform.^17^ This allele replaces Esc protein with GAL4 from the *esc* locus, by virtue of transcripts splicing into an intronic CRISPR-inserted swappable integration cassette (SIC), resulting in inclusion of an in-frame *T2A::GAL4* sequence into all transcripts (Figure 1).^17^ Notably, this SIC has flanking *attP* sites that allow germline swapping (using RMCE) with attB-flanked cassettes containing *T2A::esc* mimetic variants, by integrase-based transgenesis.^17^ In this way, we establish a panel of *esc* mimetic alleles that replace the *esc* gene with a variant that mimics an EED variant of interest (Table 1, Figure 2).

**Figure 2:**
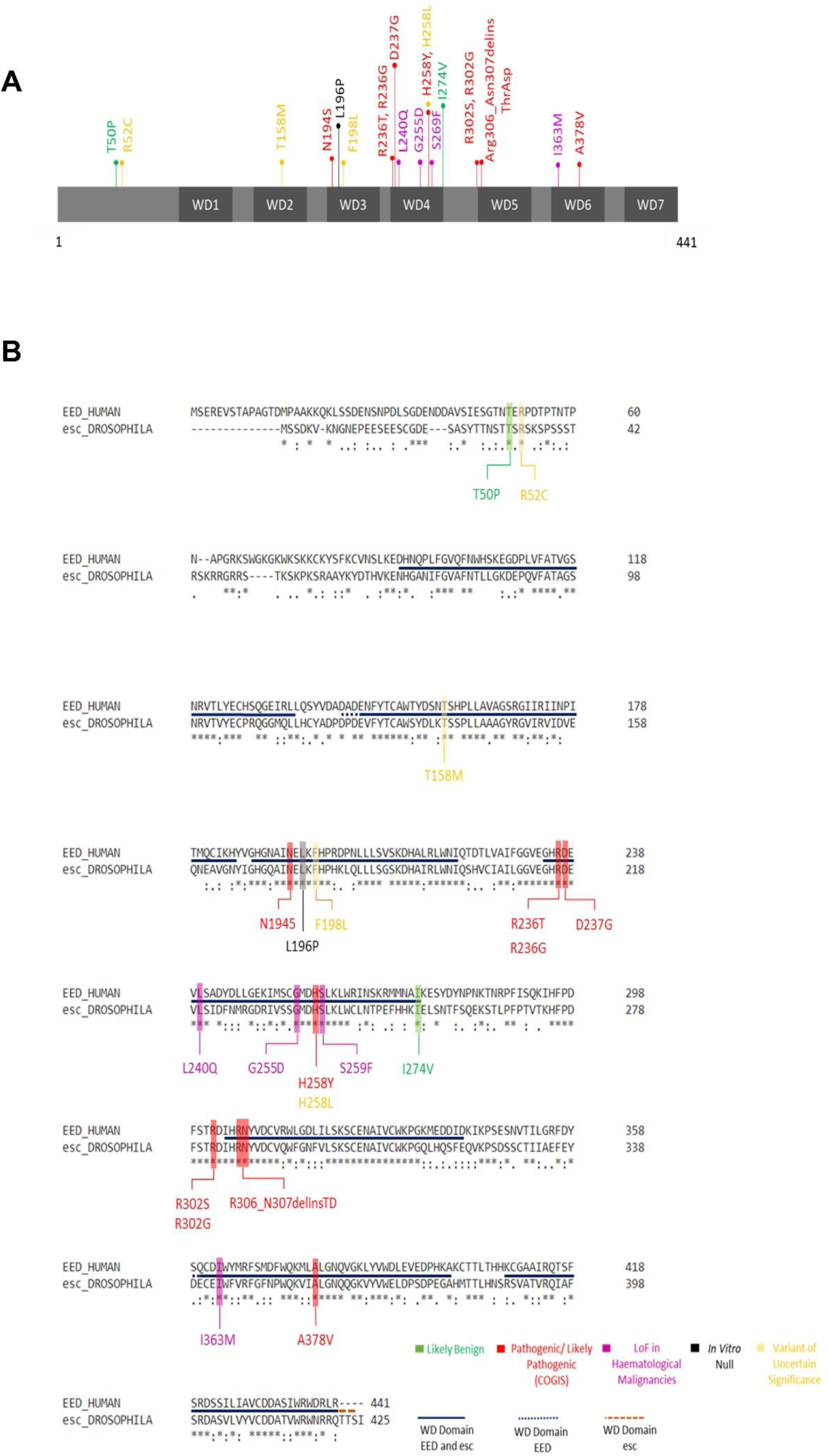
EED and Esc Protein Schematic and Alignment with EED Variants. (A) EED Protein schematic with EED variants modelled in our esc assays. (B) Protein alignment was created using Clustal Omega and FASTA files obtained from UniProt.^35–37^ UniProt IDs: EED: O75530; Esc:Q24338 ‘*’conserved residues, ‘:’conservation of amino acids of strong similar properties, ‘.’ conservation of amino acids of weak similar properties.

For the *esc* mimetic approach to be useful in functionalization assays, it is important to verify that it can phenotypically distinguish known P/LP from B/LB *EED* variants.^34^ To this end, we used RMCE to replace the *esc::T2A::GAL4* allele with either the *esc* reference sequence (*esc::Ref*), or two LB mimetic variants catalogued in gnomAD (T50P and I274V)^19^, or four COGIS mimetic variants (N194S, R236G, H258Y, R302S). We refer to these as ‘calibrating variants’.

In humans with COGIS, P/LP *EED* variants cause neurodevelopmental phenotypes in the heterozygous state. However, in *Drosophila*, *esc* heterozygosity has no overt phenotype that is easily scored. Thus, since *esc* mutant phenotypes are seen when loss-of-function (LoF) and predicted loss-of-function (pLoF) variants are present in the homozygous, hemizygous or the compound heterozygous state,^12^ we first interrogated the ability of each mimetic variant allele to produce homozygous flies. As expected, *esc::Ref* and both of the LB mimetic variants produced viable homozygous adult flies without homeotic phenotypes, at the expected Mendelian ratios (Supporting Table 1 and Supporting Table 2). In contrast, the mimetic COGIS alleles were either homozygous lethal (R236G and R302S) or produced homozygous adults less often than the expected ratio of 36% (N194S, 2%. H258Y, 7%). The expected ratio was calculated from the percentage of homozygous *esc::R*ef control flies that emerged. Additionally, all parental flies unless otherwise stated were heterozygous for a balancer allele, that is homozygous adult lethal. This decrease in expected ratios indicated that these COGIS alleles cause developmental lethality and are LoF (Supporting Table 1 and Supporting Table 2). Amongst the few adults that did emerge for N194S and H258Y, females had no overt phenotypes, but all males had extra sex combs (shown for H258Y in Figure 3).

**Figure 3:**
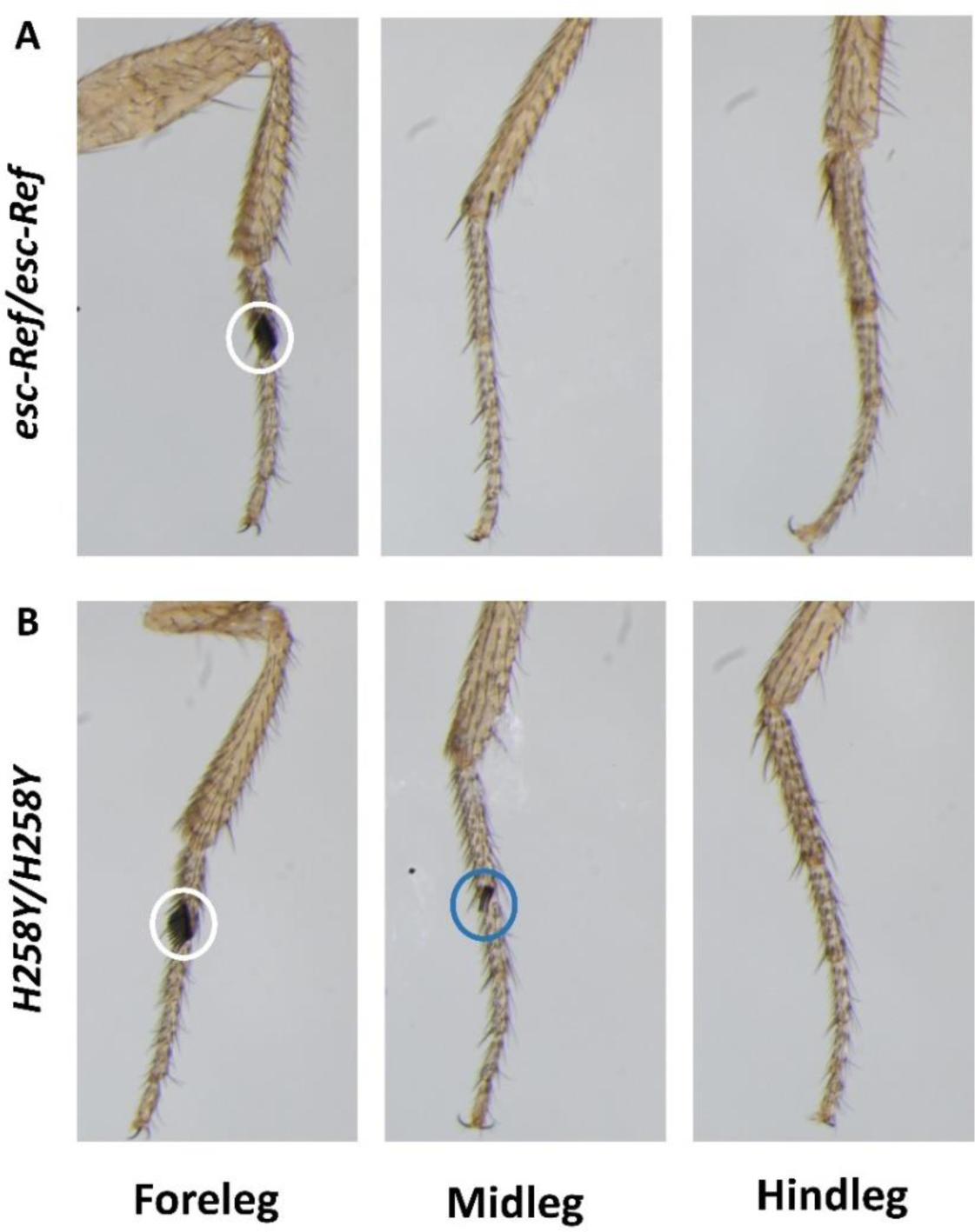
Extra Sex Comb Phenotype. Sex combs are structures seen on the first pair of legs (white circle) in males, the appearance of extra sex combs on other pairs of legs is an *esc* pLoF mutant phenotype (blue circle). **(A)** Representative image of a male fly (*esc-Ref/ esc-Ref*) with a wildtype set of extra sex combs. **(B)** Representative image of a male fly (*H258Y/H258Y*) with extra sex combs on the midleg.

We then investigated the phenotypes of flies bearing the *esc* mimetic variants (designated as *esc^#^*) in *trans* with an amorphic allele (*esc^#^/esc*^5^, *esc^#^/ Df(2L)Exel6030* or *esc^#^/esc::T2A::GAL4*). *esc*^5^ is a LoF allele on a chromosome in *cis* with *E(Pc)*^1^. ^12^ *esc::T2A::GAL4* is presumed to be a null allele.^17^ *Df(2L)Exel6030* is a deletion that removes the entire *esc* locus; ^38^ termed *esc-Df* herein. Carriers of the *esc-Df* allele are considered to be hemizygous at this locus.

We noted variable degrees of adult lethality across these genotypes (Table 2). As expected, lethality was more severe in flies with COGIS mimetic variants, particularly when the genotype included either *esc-Df* or *esc*^5^. All COGIS mimetic variants produced an extra sex comb phenotype in adult male progeny of all genotypes, whereas all LB variants produced the wildtype number of two sex combs in all genotypes (Figure 4A). We reasoned that the degree of functional impairment among variants would correlate with the severity of the extra sex comb phenotype (i.e. increased numbers of sex combs). Thus, a variant in a fly with sex combs on six legs was scored as having less function than a variant in a fly with sex combs on two legs. This framework allowed us to rank the relative function of EED variants according to severity.

**Table 2:** Observed versus Expected Frequencies of Progeny Bearing RMCE *esc* mimetic variants in the compound heterozygous or hemizygous State. LB: Likely Benign variant; C: COGIS variant; esc***^#^***: *esc* mimetic variant from column 1 in trans with esc CRIMIC-T2A-GAL4 (column 2), *Df(2L)Exel6030* (column 3) or *esc*^5^ (column 4); *Df(2L)Exel6030* is referred to as *esc*-*Df* in text; Chi-Square testing indicates that the difference in expected vs observed frequencies is significant at * p ≥ 0.05; ** p= 0.001 to 0.01; *** p= 0.0001 to 0.001; **** p < 0.0001 or not significant (ns).

|  | *esc*<sup>#</sup>/*esc* CRIMIC-T2A-GAL4 |  |  | *esc*<sup>#</sup>/*Df*(2*L*)*Exel6030* |  |  | *esc*<sup>#</sup>/*esc*<sup>5</sup> |  |  |
| --- | --- | --- | --- | --- | --- | --- | --- | --- | --- |
|  | Expected (%) | Observed (%) | Statistical Significance | Expected (%) | Observed (%) | Statistical Significance | Expected (%) | Observed (%) | Statistical Significance |
| esc-Ref | - | 75 (23%) |  | - | 200 (39%) |  | - | 72 (34%) |  |
| **T50P**<sup>LB</sup> | 61 (23%) | 74 (28%) | ns | 225(39%) | 230 (40%) | ns | 100 (34%) | 74 (25%) | ** |
| **I274V**<sup>LB</sup> | 51 (23%) | 69 (31%) | ** | 226 (39%) | 196 (34%) | ** | 101 (34%) | 85 (29%) | * |
| **N194S**<sup>C</sup> | 91 (23%) | 82 (21%) | ns | 210 (39%) | 152 (28%) | **** | 84 (34%) | 37 (15%) | **** |
| **R236G**<sup>C</sup> | 50 (23%) | 47 (21%) | ns | 197 (39%) | 155 (31%) | **** | 74 (34%) | 37 (17%) | **** |
| **H258Y**<sup>C</sup> | 72 (23%) | 59 (19%) | ns | 217 (39%) | 188 (34%) | ** | 84 (34%) | 56 (23%) | *** |
| **R302S**<sup>C</sup> | 90 (23%) | 73 (19%) | * | 193 (39%) | 149 (30%) | **** | 77 (34%) | 19 (8%) | **** |

**Figure 4:**
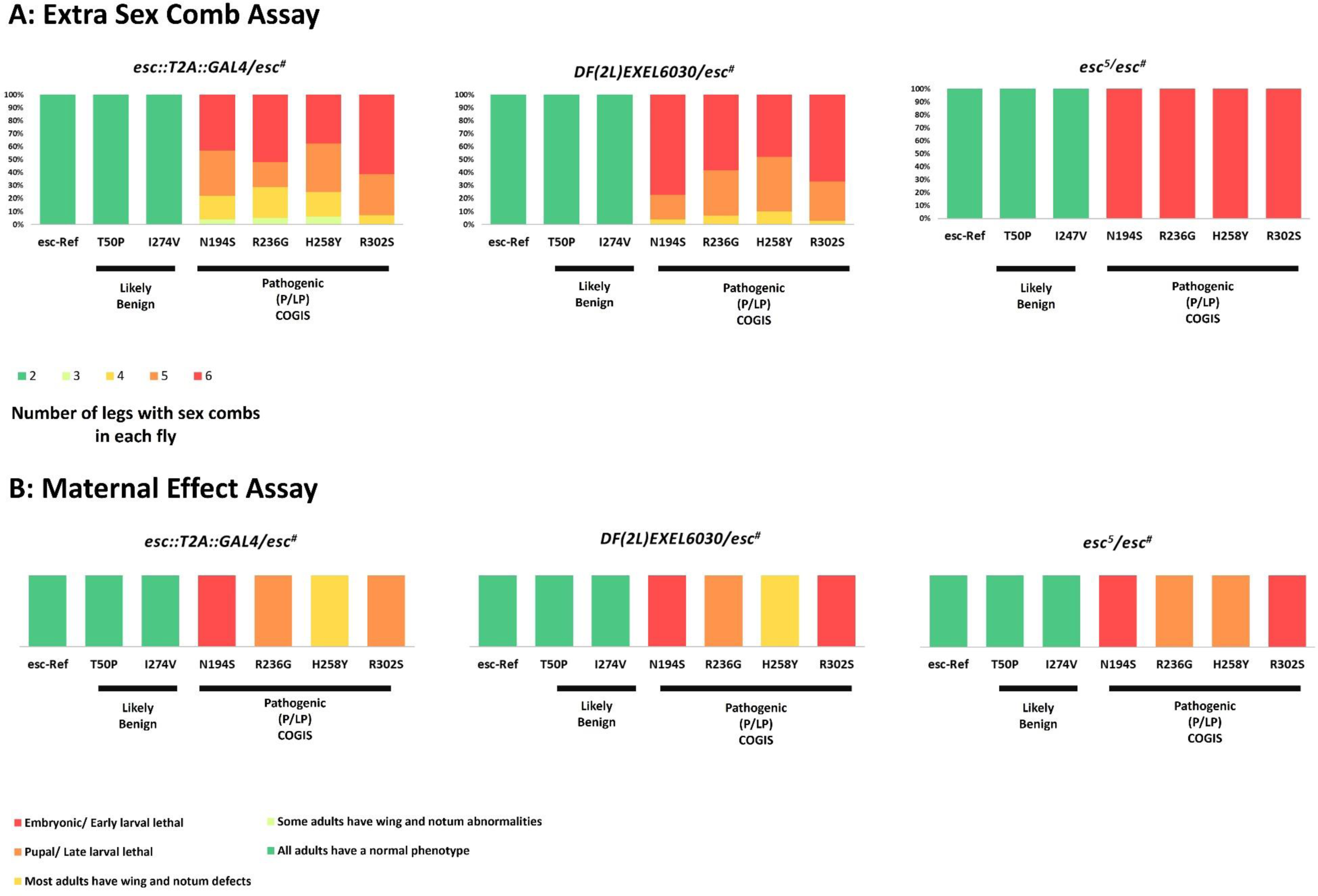
Extra sex comb and Maternal Effect Phenotypes in Calibrating Variants. **(A)** Extra sex comb phenotype in males across RMCE *esc* mimetic genotypes. **(B)** Maternal effect phenotypes across RMCE *Esc mimetic* genotypes. Each variant was assessed based on the latest developmental stage reached. esc^#^: *esc* mimetic variant; LB: Likely Benign; P:Pathogenic; LP: Likely Pathogenic Df(2L)Exel6030 is referred to as *esc-deficiency/ Df* in text

The most severe phenotype was found among flies with a COGIS variant over *esc*^5^ (*esc^#^/esc*^5^) which invariably had sex combs on all six legs. This precluded ranking of relative severity across multiple variants when they were paired with the severe *esc*^5^ allele. We posit that the increased severity of the extra sex comb phenotype in this genotype is due to the presence of the *E(Pc)*^1^ allele in *cis-* with *esc*^5^, which enhances polycomb phenotypes.^39^ However, by phenotypic scoring of male flies bearing *esc^#^/esc-Df* or *esc^#^/esc::T2A::GAL4*, each of the *esc^#^* variants could be ranked from weakest (*H258Y*) to strongest alleles (*N194S or R302S*).

We next investigated the maternal effect of each *esc* mimetic variant. Female compound heterozygotes and hemizygotes had no overt phenotype themselves. This allowed us to investigate the maternal effect of *esc* mimetic alleles on progeny. In *Drosophila*, many genes (including those in the Polycomb group) exhibit ‘maternal effects’, whereby the maternal contribution of mRNA or protein influences zygotic phenotypes.^40^ For PRC2 members including *esc*, transcripts are “maternally loaded” into developing oocytes. This is critical for early development prior to the onset of gene expression from zygotic nuclei.^41^ The progeny of *esc* null females fail to develop beyond early larval stages, even if the zygotes themselves are heterozygous for paternally-inherited wildtype *esc*.^12,41^ Thus, paternally-derived zygotic *esc* cannot fully compensate for the absence of maternally-derived *esc*. Put another way, a critical developmental window exists in the early zygote wherein maternally-supplied Esc is essential.

To test the maternal effect of each *esc* mimetic variant, we crossed mothers of genotypes *esc^#^/esc*^5^, *esc^#^/esc-Df*, and *esc^#^/esc::T2A::GAL4* to males of genotype *w*^1118^ (wildtype for *esc*). In this context, any maternally-derived Esc protein function would arise solely from the mimetic variant allele (*esc^#^*). Progeny from mothers with esc-Ref or LB variants (T50P, I274V) were phenotypically normal. Most progeny from mothers with COGIS variants died at either early to mid-larval stages or mid-larval to pupal stages, with N194S exhibiting the most severe phenotype across all genotypes; indicating the strongest loss of function. Progeny from *H258Y* mothers had the least severe phenotype; indeed we even noted that a number of adults emerged of which up to 30% had wing and/or notum abnormalities (Figure 4B and Figure 5). These data indicate that LB variants have full maternal function, while all COGIS variants show a range of loss of function.

**Figure 5:**
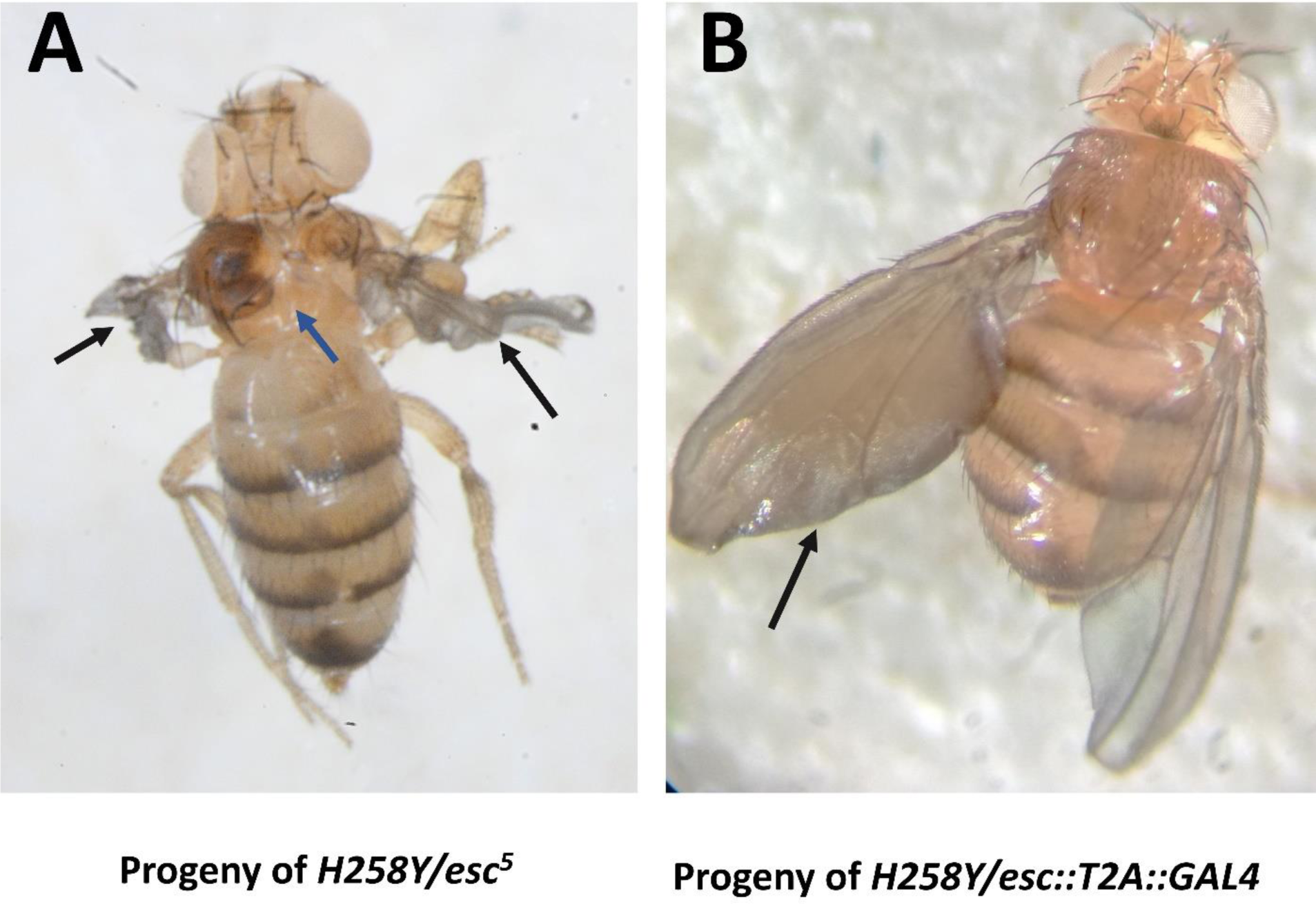
Maternal Effect Phenotypes: Representative Wing and Notum Abnormalities of Adult Progeny in Calibrating Variants. **(A)** Progeny of *H258Y/esc*^5^ (pharate adult) with wing and notum abnormalities. **(B)** Progeny of *H258Y/esc::T2A::GAL4* with one fluid-filled wing. White arrow: wing abnormalities; Blue arrow: notum abnormalities.

While scoring the phenotypes (extra sex combs and maternal effects) of the *esc* mimetic and *esc-Ref* RMCE lines, we found a variable tergite defect phenotype in adult progeny of mothers with the *esc:*:*T2A::GAL4* allele (Supporting Figure 1). However, as this phenotype was also seen in *esc:*:*T2A::GAL4* heterozygotes, including the original stock, we postulate that this phenotype is attributable to this CRIMIC insertion at the *esc* locus. Thus, we did not score this phenotype when assessing phenotypic effects on progeny.

### COGIS and Haematological Malignancy Variants are all LoF

As the RMCE approach successfully distinguished known LB and LP/P variants, we investigated an additional 14 variants, termed ‘test variants’. These included additional LP/P COGIS variants, somatic LoF haematological malignancy variants, as well as four VoUS and two component variants derived from the 306R_307NdelinsTD mutation. Additionally, we tested a known null variant L196P, an *Eed* allele observed after ethylnitrosourea-induced mutagenesis in mice.^32,33^ We chose to cross these variants to the *esc-Df* stock for subsequent assays to avoid confounders that arose using the other alleles. As noted previously, tergite phenotypes were associated with the *esc::T2A::GAL4* allele, and the *E(Pc)*^1^ allele in *cis* with *esc*^5^ enhanced the extra sex comb phenotype; thereby reducing our ability to discern differences in allelic strength between variants.

The mimetic variant of the null L196P variant was pupal lethal over *esc-Df* (Supporting Table 3); however, one single adult male did emerge with sex combs on 6 legs. We set this allele’s phenotype as the bar for complete LoF of a mimetic variant; pupal lethality and 6 sex combs. For all COGIS or haematological malignancy test mimetic variants over *esc-Df*, adult progeny did emerge, albeit often at a frequency lower than expected indicating variable degrees of developmental lethality (Supporting Table 3 and Supporting Table 4). Moreover, an extra sex comb phenotype in males was seen among all COGIS and haematological malignancy variants (Figure 6A). Similarly, based on the severity of the extra sex comb phenotype, we could discern relative allelic strength among the COGIS and haematological malignancy variants. The COGIS (calibrating and testing) and haematological malignancy variants, with the most significant effects (“highest allelic strength”) included, from strongest to weakest were: G255D, R302G, N194S, R302S, R236G, R236T, H258Y and 306R_307NdelinsTD. The latter 3 had similar allelic strengths. For the other variants, 5-44% of males had a normal sex comb number, and the remaining males had no more than 4 sex combs. These included, from strongest to weakest: A378V, D237G, S259F and I363M.

**Figure 6:**
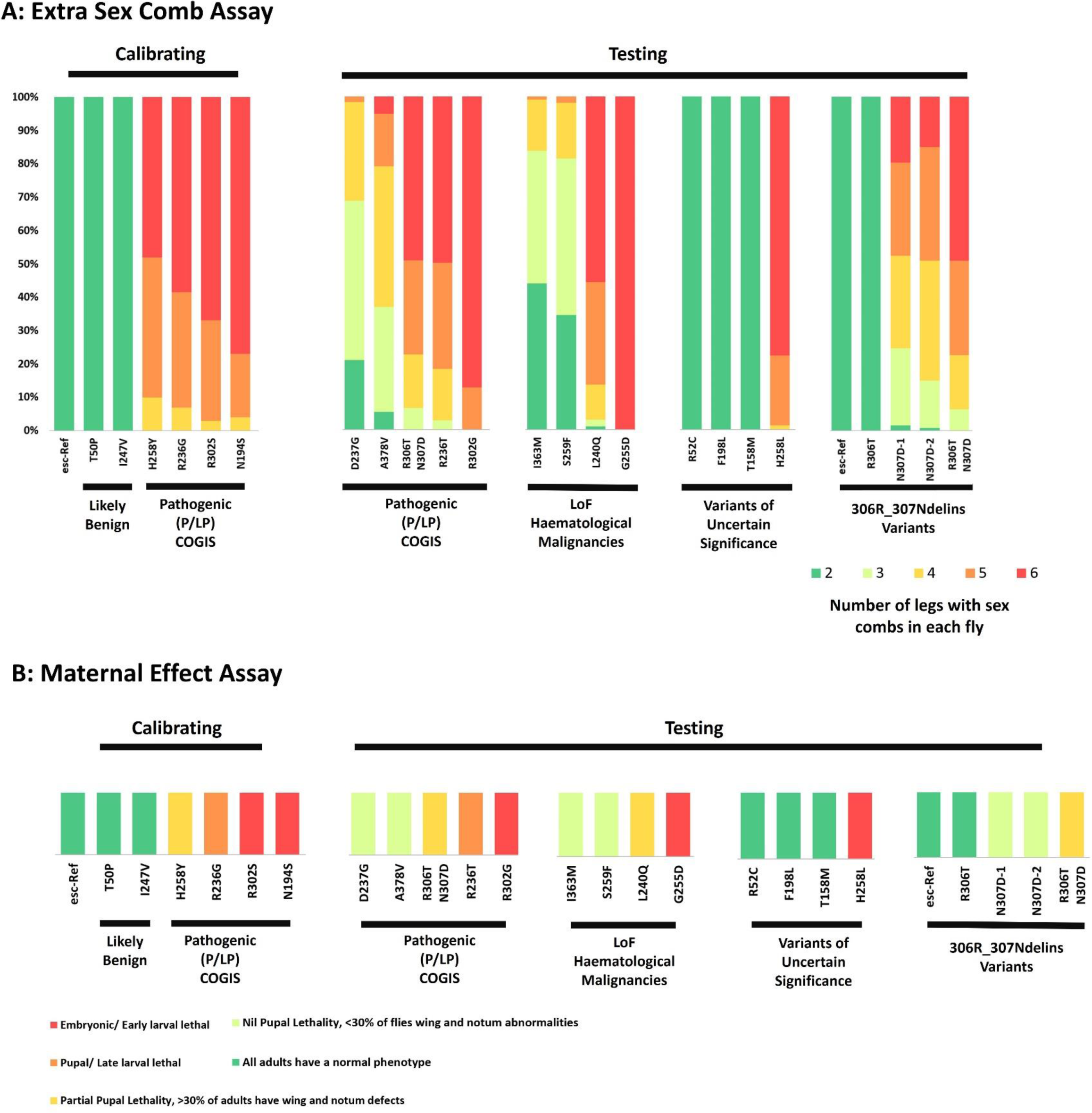
Extra sex comb and Maternal Effect Phenotypes in Testing Variants. **(A)** Extra sex comb phenotype across RMCE *esc* mimetic genotypes **(B)** Maternal effect phenotypes across RMCE *esc* mimetic genotypes. Calibrating Results are a duplicate of those seen in Figure 4. esc^#^: *esc* mimetic variant; LB: Likely Benign; P: Pathogenic; LP: Likely Pathogenic

The only variant that generated six sex combs in all male progeny was the haematological malignancy variant, G255D. Though more data are needed on COGIS patients, this observation may suggest that strong loss-of-function variants may only be observable in humans as somatic mutations. Overall, our findings emphasise that these EED variants are LoF and classifiable by the assay as LP.

We next examined maternal effects for COGIS and haematological malignancy-derived mimetic variants (Figure 6B and Supporting Table 5, Figure 7). First, progeny of mothers with variants N194S, R302S, R302G and G255D all died as embryos or as early larvae. Second, progeny of mothers with variants R236G and R236T died as late larval or pupae. For all other variants, we did observe adult progeny emerging at varying frequency; for these we further quantified the percentage with wing and/or notum abnormalities; from strongest to weakest phenotype: H258Y (65%), L240Q (40%), 306R_307NdelinsTD (34%), A378V (22%), S259F (19%), N307D (16%), D237G (6%) and I363M (4%). These results further emphasise that our approach can effectively determine the relative function of *EED* variants.

**Figure 7:**
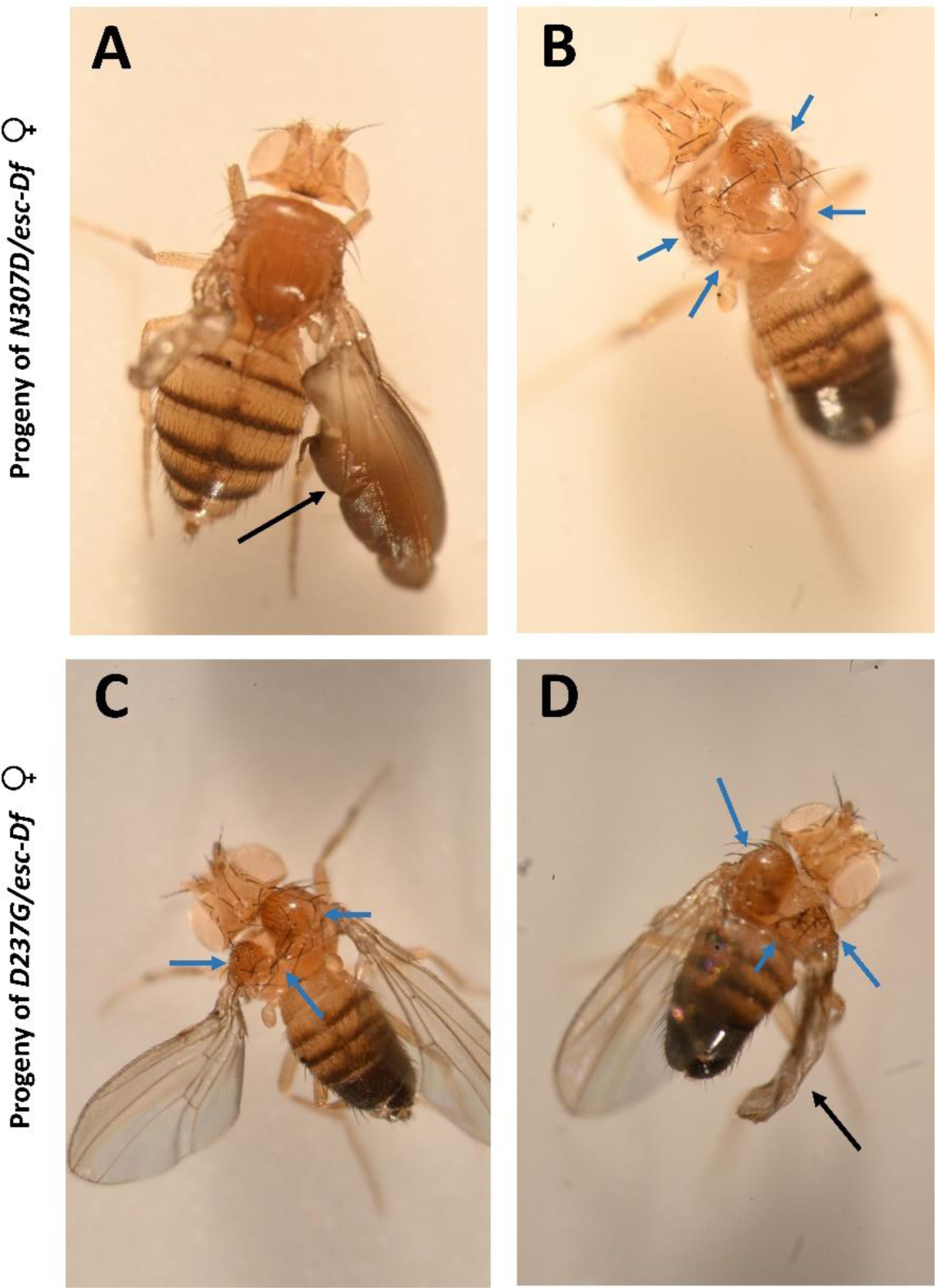
Maternal Effect Phenotypes: Representative Wing and Notum Abnormalities of Adult Progeny of Females with the Indicated Genotypes. **(A-B)** Progeny of *N307D/esc-Df* mothers: wing abnormalities seen in **(A)**, and notum abnormalities seen in **(B)** (wings removed manually for imaging purposes). **(C-D)** Progeny of *D237G/esc-Df*: notum abnormalities seen in **(C)**, and notum and wing abnormalities seen in **(D).** Black arrows: wing abnormalities; Blue arrows: notum abnormalities

### Reclassification of four *EED* VoUS

We next tested four VoUS, R52C, T158M, FI98L and H258L over *esc-Df* to aid in their reclassification (Table 1). All except H258L proved to have wildtype function, which we reclassify here as likely benign. Flies hemizygous for R52C, T158M and FI98L were viable (Supporting Table 3), all adult male progeny had normal sex comb numbers, and no maternal effects were observed in progeny of hemizygous mothers (Figure 6 and Supporting Table 5). In contrast, while hemizygous H258L adults were fully viable, all adult male progeny had extra sex combs (Figure 6A) and all progeny of hemizygous mothers died at the embryonic to early larval stages (Figure 6B). These results indicate that R52C, T158M and FI98L are LB variants while H258L is LoF and would be classified as a LP variant by this assay.

### Pathogenicity of *306R_307NdelinsTDR306T* arises mainly from the *N307D* component

The COGIS indel *306R_307NdelinsTD* alters two adjacent amino acid residues. We tested which amino acid change(s) contribute to pathogenicity. We generated independent *R306T* and *N307D* mimetic alleles. Extra sex combs were found on hemizygous N307D but not R306T males (Figure 6A). Similarly, only progeny of *N307D/esc-Df* exhibited any maternal effect, as observed by the number of adult progeny that had wing and/or notum phenotypes (Figure 6B And Supporting Table 5). Interestingly, the LoF impact of 306R_307NdelinsTD was worse than N307D alone. 49% of male 306R_307NdelinsTD hemizygotes had six sex combs, compared to 15-20% of *N307D* hemizygotes (Figure 6A). Similarly, progeny of *306R_307NdelinsTD/esc-Df* females had 38% wing and/or notum defects, compared to 16-23% seen among progeny of *N307D/esc-Df* females (Figure 6B and Supporting Table 5). These data show that the low cost and high efficiency of the *Drosophila* approach offers a means of testing how each amino acid change contributes to pathogenicity.

### The SIC in *esc* caused male sterility among homozygotes and compound heterozygotes

We observed that adult males homozygous or hemizygous for mimetic variants modelled using the RMCE approach produced very few progeny, or failed to produce progeny at all. To investigate this male infertility, we crossed compound heterozygous and hemizygous males from our calibrating set of variants (*esc^#^/ esc*^5^, *esc-Df* or *esc::T2A::GAL4*) to *w*^1118^ females. We found that males with any mimetic LB or LP variant in *trans* with the *esc-Df* or *esc::T2A::GAL4* alleles were infertile. In contrast, males with any LB or LP calibrating mimetic variant allele over the *esc*^5^ allele were fertile. Relevant to these findings is that the [-] strand directed *esc* is intronic to the larger [–] strand directed *Pde1c* gene, which is male fertility-defective when mutant.^42^ The *esc-Df* allele deletes *Pde1c* in its entirety,^42^ and it is possible that the splice acceptor within the artificial exon created by the CRIMIC insertion of *esc::T2A::GAL4* forces aberrant splicing of *Pde1c*, resulting in a loss of function allele. Thus, a parsimonious explanation for the selective male infertility of variants over *esc-Df* or *esc::T2A::GAL4,* but not over *esc*^5^, is a loss of *Pde1c* function.

### Expression of pathogenic *UASz-esc* mimetic variants causes lethality

In a second approach based on the CRIMIC strategy, we used the *esc::T2A::GAL4* line to drive expression of *UASz-esc* mimetic variants in the same spatial and temporal pattern as endogenous *esc*, so we could examine rescue of *esc* null phenotypes. By ϕC31 integrase transgenesis, we generated nine lines of flies, each with a different *UASz-esc* mimetic variant integrated into *attP2*. We selected five COGIS variants, two LB variants and the two individual missense components of the *306R_307NdelinsTD* indel.

Expression of COGIS variants (N194S, R236G, H258Y, R302S and 306R_307NdelinsTD) caused lethality prior to adulthood, whereas expression of LB variants (T50P and I274V) resulted in viable adults at the expected Mendelian ratio (Table 3). We tested the individual amino acid changes comprising 306R_307NdelinsTD. Expression of R306T resulted in viable adults but expression of N307D reduced adult viability (fewer than 50% of expected numbers) (Table 3) and all males (3) had extra sex combs. These results are consistent with those seen from the RMCE strategy. However, in this assay expression of all COGIS variants produced lethality which precluded ranking variants by allelic strength by adult viability alone.

**Table 3:** Observed and Expected Frequencies of *UASz-esc* mimetic Variant Expression. *esc^#^: esc* mimetic variant from column 1. Chi-Square testing indicates that the difference in expected vs observed frequencies is: ns: not significant or significant: * p ≥ 0.05; ** p= 0.001 to 0.01; *** p= 0.0001 to 0.001; **** p < 0.0001. *attP2*: negative control (nil transgene expression)

| Line | *esc::T2A::GAL4* /+; *UASz-esc*<sup>#</sup>/+ |  |  |
| --- | --- | --- | --- |
|  | Expected (%) | Observed (%) | Statistical Significance |
| **Controls** |  |  |  |
| *esc*-Ref | 19 (50%) | 19 (50%) | ns |
| *attP2* | 25 (50%) | 25 (50%) | ns |
| **Likely Benign Variants** |  |  |  |
| T50P | 22 (50%) | 26 (59%) | ns |
| I274V | 34 (50%) | 37 (54%) | ns |
| **Likely Pathogenic/ Pathogenic COGIS Variants** |  |  |  |
| N194S | 7 (50%) | 0 (0%) | *** |
| R236G | 10 (50%) | 0 (0%) | **** |
| H258Y | 11 (50%) | 0 (0%) | **** |
| R302S | 9 (50%) | 0 (0%) | **** |
| 306R_307NdelinsTD | 39 (50%) | 0 (0%) | **** |
| **306R_307NdelinsTD Component Variants** |  |  |  |
| R306T | 27 (50%) | 30 (56%) | ns |
| N307D | 78 (50%) | 18 (23%) | **** |

### Expression of LB *UASz-esc* mimetic variants rescues *esc* lethality

Next, we investigated the ability of *UASz-esc* mimetic variants to rescue a lethal *esc* phenotype. We noted from the above experiment that overexpression of LP variants caused lethality, but we wanted to see whether a rescue-based assay might yield additional data that would better subcategorize allelic strength between the *UASz-esc* mimetic variants.

The genotype *esc::T2A::GAL4 /esc-Df* is adult lethal. We crossed *esc::T2A::GAL4/CyO* females to *esc-Df/CyO ; UASz-esc^#^*males bearing a variety of test *esc^#^* alleles. We then screened for the following genotype: *esc::T2A::GAL4 /esc-Df ; UASz-esc#/+*. The presence of progeny with this genotype indicated that the expression of the *esc^#^* mimetic variant under the control of the endogenous *esc* promoter via the GAL4-UAS system rescued lethality.

First, we observed full rescue of *esc* lethality, with no observable phenotypes, (Table 4) by *UASz-esc-Ref*, all LB *UASz-esc^#^* variants and also *UASz-R306T* (Table 4). In contrast, all COGIS *UASz-esc^#^* mimetic variants failed to rescue *esc* lethality (Table 4). *UASz-N307D* exhibited 1% rescue of lethality, and the few emerging adults had wing and notum defects, and the sole male had sex combs on all six legs.

**Table 4:** Rescue of a Lethal Phenotype: Observed and Expected Frequencies of *UASz-esc* Mimetic Variants. *esc*^#^: *esc* mimetic variant from column 1. Chi-Square testing indicates that the difference in expected vs observed frequencies is: ns: not significant or significant: * p ≥ 0.05; ** p= 0.001 to 0.01; *** p= 0.0001 to 0.001; **** p < 0.0001. *attP2*: negative control (no transgene expression)

| Line | *esc::T2A::GAL4 / Df(2L)EXEL6030; UASz-esc<sup>#</sup>/+* |  |  |
| --- | --- | --- | --- |
|  | Expected (%) | Observed (%) | Statistical Significance |
| **Controls** |  |  |  |
| esc-Ref | - | 215 (35%) | - |
| *attP2* | 0 (0%) | 0 (0%) | - |
| **Likely benign Variants** |  |  |  |
| T50P | 202 (35%) | 217 (38%) | ns |
| I274V | 196 (35%) | 215 (38%) | ns |
| **Likely Pathogenic/ Pathogenic COGIS Variants** |  |  |  |
| N194S | 88 (35%) | 0 (0%) | **** |
| R236G | 90 (35%) | 0 (0%) | **** |
| H258Y | 60 (35%) | 0 (0%) | **** |
| R302S | 82 (35%) | 0 (0%) | **** |
| 306R 307NdelinsTD | 67 (35%) | 0 (0%) | **** |
| **306R_307NdelinsTD Component Variants** |  |  |  |
| R306T | 212 (35%) | 221 (36%) | ns |
| N307D | 89 (35%) | 3 (1%) | **** |

As we obtained *esc::T2A::GAL4 /esc-Df; UASz-esc^#^/+* females from both LB variants and also *UASz-R306T* and *UASz-N307D*, we tested if these variants could rescue the lethal maternal effect found in progeny of *esc::T2A::GAL4 /esc-Df* mothers. We crossed *esc-Df/CyO* males to *esc::T2A::GAL4 /esc-Df; UASz-esc^#^/+* females. We hypothesised that the expression of LB variants and *UASz-R306T* in females (*esc::T2A::GAL4 /esc-Df ; UASz-esc^#^/+)* would not have an observable effect on progeny, and moreover that a maternal copy of a benign *UASz-esc^#^* transgene would rescue the emergence of adult progeny that were *esc::T2A::GAL4 /esc-Df : +/+,* with no *UASz-esc^#^* transgene. As expected, adult progeny that were mostly phenotypically normal emerged from hemizygous mothers with *esc-Ref* or T50P, I274V and R306T variants (Table 5). Additionally we were able to recover female and male progeny that were *esc::T2A::GAL4 /esc-Df* ; +/+. However, all adult male progeny of genotype *esc::T2A::GAL4 /esc-Df* ; +/+ had extra sex combs. In contrast, we found that a maternal supply of *UASz-N307D* was unable to rescue the emergence of any adult *esc::T2A::GAL4 /esc-Df* : +/+ progeny. Moreover, all adult progeny from this cross carried the CyO balancer and also exhibited wing or tergite abnormalities (Table 5) Thus, a maternal supply of *UASz-N307D* could not provide sufficient Esc function in the absence of a zygotic *esc* wild-type allele, emphasizing the LoF effects of this variant. This further demonstrates that the LB variants chosen in this study have *esc* activity similar to that of *esc-Ref* and are most likely truly Benign variants. Unfortunately, this experiment was not feasible for COGIS variants, because females with the required genotype were adult lethal. Such an assay might still useful as a second tier to further demonstrate that a variant of interest has Esc activity similar to that of the *esc-Ref* or for variants with subtler LoF effects such as N307D.

**Table 5:** *UASz-esc^#^* Likely Benign Mimetic Variants Rescued Lethality of Combined Maternal and Zygotic *esc* Nulls. *esc^#^*: esc mimetic variant from column 1

|  | Total Adult Progeny | *esc::T2A::GAL4 / DF(2L)EXEL6030; UASz-esc<sup>#</sup>/+* | *esc::T2A::GAL4 / DF(2L)EXEL6030* |
| --- | --- | --- | --- |
| **Controls** |  |  |  |
| esc-Ref | 53 | 16 | 3 |
| **Likely benign Variants** |  |  |  |
| T50P | 118 | 10 | 5 |
| I274V | 66 | 21 | 2 |
| **306R_307NdelinsTD Component Variants** |  |  |  |
| R306T | 117 | 25 | 5 |
| N307D | 7 | 0 | 0 |

### A knock-in *H258Y* allele shows partial loss-of-function effects in male and female flies

The third approach we employed was homology-directed repair (HDR) by CRISPR/Cas9. We made an *esc* mimetic allele of the *EED H258Y* COGIS variant (*H258Y^KI^*) (Supporting Figure 2). We also made a control ‘reference sequence’ allele (*esc-Ref^KI^*) to control for the single nucleotide PAM sequence mutation at each of the two gRNA target sequences used in the HDR strategy.

We recovered four independently-generated crispant lines of the *H258Y^KI^* mimetic allele and five independently-generated crispant lines of the *esc-Ref^KI^* allele. These were all outcrossed in a *w*^1118^ background for six generations to remove second site mutations. Because the *esc-Ref^KI^* crispant lines were entirely wildtype, we believe that the cassette had little to no impact on *esc* function. Additionally, we removed this cassette from one line each of the *esc-Ref^KI^* and *H258Y^KI^* lines, and observed similar results as we report below.

We tested *H258Y^KI^* in the homozygous, compound heterozygous and hemizygous states. The four *H258Y^KI^* lines were homozygous lethal. This was not due to the PAM site mutations; all *esc-Ref^KI^* lines were homozygous viable with no observable phenotype. We next tested these alleles in *trans* with LoF alleles *esc*^5^, *esc-Df* and *esc::T2A::GAL4*. We found that all *esc-Ref^KI^* lines were viable and fertile over these three null alleles (Supporting Table 6 and Supporting Table 7). However, all *H258Y^KI^* lines demonstrated some lethality over two of three alleles. Flies with genotype *H258Y^KI^/esc::T2A::GAL4* made it to adulthood at the expected Mendelian ratios, but 69/71 of these males had an extra sex comb phenotype (Supporting Table 6 and Supporting Table 7). In contrast, both *H258Y/esc*^5^ and *H258Y/esc-Df* genotypes made it to adulthood at reduced rates of 13% and 11-26%, respectively; all male progeny of these genotypes had extra sex combs (Figure 8). As expected, this extra sex comb phenotype was not seen in any of the genotypes of *esc-Ref^KI^* males.

**Figure 8:**
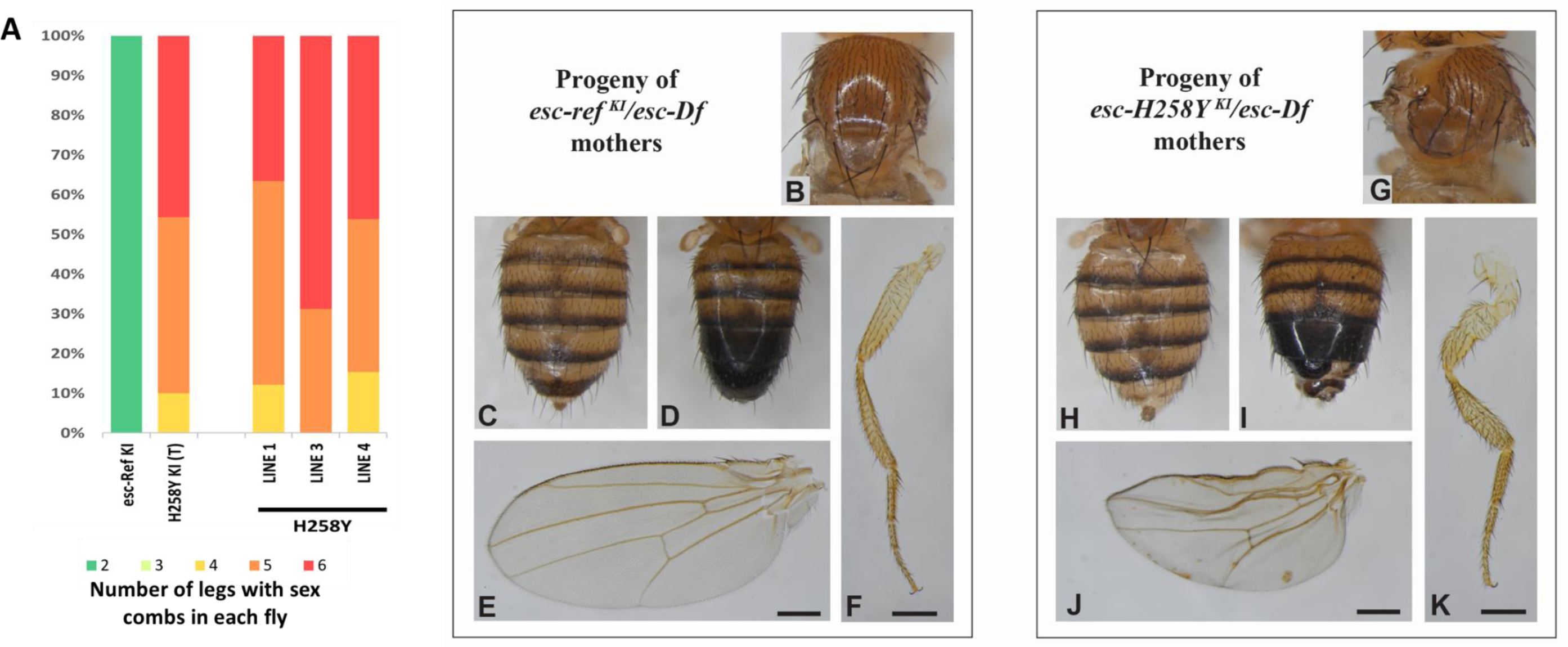
Extra Sex comb Phenotype in *H258Y*^KI^ Males and Maternal Effect Phenotypes of *H258Y ^KI^* in adult progeny. **(A)** Genotype of males are *H258Y^KI^/esc-Df* and *esc-Ref^KI^/esc-Df.* H258Y^KI^ (T) represents the combined number of males across lines 1, 3 and 4. **(B-F)** Progeny of *esc-ref ^KI^/esc-Df* females crossed with *w*^1118^ males had no phenotypes in notum **(B)**, abdomen **(C-D)**, wings **(E)** and legs **(F)**. **(G-K)** Progeny of *esc-H258Y ^KI^/esc-Df* mothers and *w*^1118^ fathers exhibited a variety of phenotypes in the notum (**G**) abdominal segment A6 (**H-I**), wing **(J),** and/or legs (**K**). Scale bars equal 250 μm.

In females, *H258Y^KI^* compound heterozygotes and hemizygotes had no overt phenotype, so we investigated maternal effects. First, we examined the progeny of females of genotype *esc-Ref ^KI^*/*esc-Df*., crossed to *w*^1118^ males (wildtype for *esc*). This progeny emerged as adults with no overt phenotype, and at expected ratios indicating an absence of any developmental lethality (Figure 8 B-F and Table 6). In contrast, progeny of *H258Y ^KI^*/*esc-Df* females and *w*^1118^ males were mostly pupal lethal. The few progeny that emerged as adults all exhibited a range of phenotypes. All females and most males had tergite pigmentation defects of the A6 segment (Figure 8 H-I and Table 6). Half of those flies also had wing phenotypes, and in some cases, we also observed notum and/or leg phenotypes (Figure 8 G, J and K). The spectrum of wing phenotypes was variable, ranging from wings with mild defects in shape and veins pattering to more severe as blistered or unexpanded wings. Similarly, the progeny of *H258Y^KI^*/*esc*^5^ females crossed with *w*^1118^ males presented the same malformation phenotypes as described above. (Figure 8). Together, the extra sex comb and maternal effect phenotypes examined above both indicate that the *H258Y* crsipant allele is LoF. These experiments demonstrate that a *Drosophila* knock-in model is another approach that can interrogate LoF effects in *EED* variants.

**Table 6:** Maternal effect phenotypes of *H258Y^KI^* in adult progeny. Females of genotypes indicated were crossed to *w*^1118^ males. Adult progeny were scored for maternal effect phenotype, with penetrance indicated by the number (and %) of progeny with each phenotype. Data were taken from a single KI line for each genotype.

| Progeny of mothers<br>*esc-ref*<sup>KI</sup>/*esc-Df* |  |  | Progeny of mothers<br>*esc-H258Y*<sup>KI</sup>/*esc-Df* |  |
| --- | --- | --- | --- | --- |
| Females (%) | Males (%) | Phenotype | Females (%) | Males (%) |
| 59 (100%) | 30 (100%) | No phenotype | 0 | 1 (9%) |
| 0 | 0 | Only wing | 0 | 1 (9%) |
| 0 | 0 | Only abdominal segment | 9 (50%) | 3 (27%) |
| 0 | 0 | Wings, abdominal segment | 7 (39%) | 1 (9%) |
| 0 | 0 | Wings, abdominal segment, notum | 1 (6%) | 1 (9%) |
| 0 | 0 | Wings, abdominal segment, notum, leg | 1 (6%) | 2 (18%) |
| - | 0 | Extra sex comb, abdominal segment | - | 2 (18%) |
| 59 | 30 | TOTAL | 18 | 11 |

## Discussion

We demonstrate that assaying the function of mimetics of *EED* variants in *Drosophila* is an effective strategy to interrogate variant function with clinical predictive value. We assayed simple, canonical phenotypes resulting from altered PRC2 activity. These canonical phenotypes included extra sex combs, other developmental abnormalities and lethality resulting from maternal effects. We investigated variant function in multiple zygotic and maternal genotypes across 22 *EED* variants, generating additional and in some instances novel *in vivo* functional data on LP COGIS variants, LoF haematological malignancy variants, and VoUS. In these VoUS that were found in patients, our functional data can contribute to the updating of their classification.

We wished to test the utility and consistency of numerous technical strategies for *EED* variant assessment. All *esc* mimetic assays developed here are convenient for discriminating variants for LB or LoF activity. Moreover, we found that the severity of the fly phenotype predicted the severity of the allele, and data on severity is consistent across all assays to the extent that a variant in the extra sex comb assay predicts the severity of the same variant in the maternal effect assay. For instance, variants that produced more than 50% of males with sex combs on all six legs were likely to exhibit larval or pupal lethality in the maternal effect assay. Similarly, variants that produced males that had sex combs on fewer than 6 legs, or when fewer than 50% of males had sex combs on 6 legs, resulted in zygotes that were likely to reach the adult stage in the maternal effect assay.

As all strategies proved similarly capable of discriminating LB and LoF variants, we find that each strategy can be exploited for its specific advantages, depending on the goal. The least expensive and fastest option is overexpression of the *UASz-esc^#^* using *esc::T2A::GAL4.* This provides a simple yes / no answer to whether an *EED* variant is LoF/LP. We envision that this would be sufficient for quick classification of a rare variant of concern found in the clinic, such as in a patient with neurodevelopmental delays with a VoUS that could not be proven to be *de novo* because parental samples were unavailable. Also, when diagnostically-suspicious VoUS are present in more than one gene, an assay of this type could discern causality attributable to the *EED* variant. However, this particular assay does not permit quantitative assessment of the allelic strength of a variant. This would require the more time-consuming approach of rescuing *esc::T2A::GAL4/ esc-Df* nulls with *UASz-esc^#^*, as it requires building *UAS-transgenes* into an *esc-Df* background. Yet, this assay would use the same *UASz-esc^#^* transgene as the simple overexpression assay, so could be applied as a follow up test. Finally, swapping *T2A::esc#* for the *esc::T2A::GAL4* cassette (the RMCE approach) offers the potential for high resolution data on the nature of the allele. Importantly, the *esc* mimetic variant is expressed at near physiological levels, in contrast to GAL4/UAS-driven overexpression of variants. Moreover, *esc^#^* alleles can be placed in *trans* with any other *esc* allele of interest to examine patient bi-allelism and better infer *esc^#^* disease mechanism to test if alleles are antimorphic, neomorphic or hypermorphic.

Our data showed that three of the four VoUS tested were LB, whereas H258L was found to be LoF. The T158M VoUS has an allele count of 3 in gnomAD V2.1.1 (July 21^st^ 2023).^19^ Since recruitment of adults to gnomAD selects somewhat against patients with childhood-onset neurodevelopmental syndromes, and there are only 12 reported individuals with COGIS to date,^22^ this T158M variant had a reasonable prior probability of being LB. In contrast, classification of the EED R52C VoUS was less clear. It was reported in an individual suspected to have hereditary breast and ovarian cancer.^23^ Previous literature had not implicated LoF *EED* variants in this cancer syndrome, and the degree to which this specific *EED* variant was thought to be contributing to the phenotype of the patient (as compared to other somatic and germline variants they may have had) was not annotated in the ClinVar database (July 21^st^ 2023). Thus, our data could find utility in ruling out this variant as a contributor to pathology. Additionally, we have limited clinical information on the patient with F198L, and thus are unaware if they have phenotypes related to COGIS. Our identification of this variant as LB may help rule this variant out as a contributor to pathology. In contrast, the individual with H258L had a clinical diagnosis of Weaver Syndrome prior to whole exome sequencing.^21^ The variant was classified as a VoUS as inheritance could not be determined; however, our results suggest that this variant has a pLoF functional impact which would be consistent with diagnosis. Additionally, another pathogenic COGIS mutation (H258Y) has been reported at this amino acid position, and is pLoF in our experiments. This emphasises that mutations at this amino acid position may result in pLoF effects and cause COGIS.

We also examined if our strategy could be used to discriminate the pathogenic amino acid change of interest in the 306R_307NdelinsTD variant. Our results demonstrate that R306T in isolation is LB, whereas N307D is LoF. This discrimination provides data pertinent to future variant classification where variants at either amino acid position may be encountered in a clinical genetic or genomic test result. We also showed that the original 306R_307NdelinsTD variant has more significant effects on the EED protein than does N307D in isolation, most likely owing to a synergistic effect of the two variants. Though of mechanistic interest, the clinical utility of teasing apart the causality of specific missense variants within in-frame indels is less clear. Nevertheless, the process would inform interpretation of future identified missense variants and potentially assist therapeutic development by illuminating which residues are critical for specific protein functions.

Functional evidence is essential in variant interpretation, as evidenced by the fact that functional evidence is allocated a rating of PS3 (Strong evidence of pathogenicity) and BS3 (Strong evidence of likely benign impact) within the ACMG-AMP criteria.^34^ These criteria specify that functional data should be generated from well-established functional assays. In 2020, the Clinical Genome Resource (ClinGen) Sequence Variant Interpretation Working Group provided recommendations for the application of the PS3 and BS3 criteria. The authors outlined that an assay should model or test pathogenesis or mechanism of disease, and include a minimum of 11 B/LB and P/LP controls (i.e. 11 in aggregate over all 4 variant subclasses) to validate the assay. Our work fulfilled these two criteria. *EED* variants are hypothesised to cause disease through a LoF mechanism. By using defined LoF phenotypes in *Drosophila* as our assay outcomes, we by proxy investigated if variants assayed had LoF effects. Additionally, we tested two LB and nine LP COGIS variants, which produced expected results in our assays. Thus, our assays can be used to generate data on *EED* VoUS to aid in interpreting variants that may be encountered in the clinical space.

The ClinGen recommendations for the application of the PS3 and BS3 criteria state that data from an assay using eleven minimum controls can be used as moderate level evidence in the ACMG-AMP variant interpretation criteria, when an Odds of Pathogenicity (OddsPath) cannot be calculated.^18^ Thus, the outcome of testing VoUS in our EED assays can be applied at moderate level evidence. Notably, the ClinGen recommendations also stated that, for functional evidence from model organisms, the strength of evidence can be adjusted based on rigour and reproducibility. We believe that these assays are robust and well-validated because our *esc* mimetic variants showed similar results when tested using three differing methodologies, and also that the known LP and LB variants faithfully recapitulated their function in humans.

There are two notable limitations in the use of our assays, anchored in the underlying biology of the system we chose. Any variant tested in the extra sex combs and maternal effect assay must occur at an amino acid position conserved between humans and *Drosophila*, which may exclude testing of a fraction of variants seen in human populations, or at the very least mandate caution in interpretation of assay results. However, all LP/P *EED* variants, encountered thus far are at identical amino acid positions between humans and *Drosophila*, and thus most clinically relevant variants may be amenable to testing through our assays. The second limitation regards our lack of interrogation of gain-of-function (GoF) effects. GoF effects may not be relevant clinically, as there are no reported *esc* or *EED* GoF alleles to date. However, variants in haematological and other malignancies operate under less stringent biological constraints than do constitutional variants, so while EED GoF variants may not be viable in the germline, there may be cellular contexts in which EED GoF variants might be observed as postzygotic somatic mutations. It is also important to note that the design of our assays intrinsically assumes that a GoF variant would manifest with a distinct phenotype distinguishable from that of the LoF function phenotypes we observed. We note here that E(z), another PRC2 component in the fly, has a documented GoF variant that has a distinct and separate phenotype from documented E(z) LoF variants.^43,44^ Additionally, a *Drosophila* mimetic assay designed to assess a rare human EZH1 variant found the variant to be GoF, in part assessed by evidence for increased H3K27 trimethylation for the mimetic variant relative to a wildtype allele of the orthologous *E(z)* gene.^11^ Therefore, if an *EED* GoF variant is assayed, we would expect the presence of some, if not all of the phenotypic features associated with the E(z) GoF variant in the resultant Esc mimetic GoF fly.

*EED* variants seen in COGIS and haematological malignancies were functionally LoF, as expected. These results match previous findings from published functional data indicating a LoF impact.^3,4,29,31,45^ Although we showed that variants had different effects on severity in the assays tested, this may not predict specific clinical phenotypes in humans with a high degree of confidence. In COGIS, different individuals with the same variant manifest variability in their clinical phenotype, and there are no variant-specific relationships between genotype and phenotype yet reported for this syndrome. This clinical variability is exemplified by the three independent individuals with the N194S variant, wherein the presence or absence of features such as structural brain abnormalities, seizures, cardiac and ocular abnormalities were not consistent amongst reported cases.^20–22^ Additionally, there is no consensus on what methodology should be used to rank severity in patients. Similar to that found with constitutional *EED* variants among patients with rare syndromes, haematological malignancies associated with *EED* missense LoF mutations do not yet have prognostic indicators based specifically on the *EED* variants.^29^ Therefore, we do not know whether an *EED* variant that had more severe LoF effects in our assays would cause more severe disease as a constitutional or somatic mutation in a living person. As mentioned above, the diagnostic utility of a test like this one is likely to be greatest for patients who have diagnostically-suspicious VoUS in *EED*, such that causality for phenotypic features could be attributed to the *EED* variant or shunted away from it. The rarity of most VoUS means that population frequencies alone will never be sufficient to classify rare variants correctly, and if *de novo* status cannot be determined due to the unavailability of parental samples, functional assays like this one may resolve the classification.

Generating *esc* mimetic mutations in the fly was a viable and successful avenue to interrogate the functional effects of human *EED* variants. This provides a basis for our new hypothesis that once suitable assays are found, this approach can be used for the other PRC2 genes; particularly *EZH2* and *SUZ12*. Germline pathogenic mutations in *EZH2* and *SUZ12* cause Weaver syndrome and Imagawa-Matsumoto syndrome respectively, while somatic mutations and dysregulation in these genes have also been implicated in malignancies. Additionally, up to 1% of the general population have a rare variant in any of these three genes (supplemental methods),^19,46–48^ and assays such as these would have utility to determine if diagnostically-suspicious variants are truly benign, or causal for phenotypic features seen in patients, aiding in the efforts to resolve the challenge of VoUS in the PRC2 genes.

## Methods and Materials

### Fly Husbandry

Fly stocks were maintained at room temperature on standard cornmeal food, and crosses were set up and maintained at 25°C unless otherwise stated. For each experiment, a minimum of 2-3 independent crosses were set up, with 2-3 replicates of each cross. An *esc* reference control was included in each experimental batch.

*Drosophila* stocks were either generated by us (below), obtained from the Bloomington *Drosophila* Stock Centre Indianna University (indicated by a BL number), or were generously donated. BL3142: *esc[5] E(Pc)[1]/SM5* (referred to as *esc*^5^), BL3623: *esc[21] b[1] cn[1]/In(2LR)Gla, wg[Gla-1]; ca[1] awd[K]* (referred to as *esc*^21^), BL79282*: y[1] w[*]; TI{GFP[3xP3.cLa]=CRIMIC.TG4.0}esc[CR00712-TG4.0] Pde1c[CR00712-TG4.0-X] /SM6a/CyO* (referred to as *esc-T2A::GAL4*), BL7513: *w*^1118^*; Df(2L)Exel6030, P{XP-U}Exel6030/CyO*, *w*^1118^ (referred to as *esc-Df*), *Adv/CyO* (generously donated by Dr. Gino Laberge (Genome Prolab Inc. (Quebec); *w*^1118^; ES/*CyO* (Balancer line); *w*^1118^ (used as control).

### Variant Selection

All variants selected were in amino acid residues that are identical between the human and *Drosophila* orthologues. LB variants in EED were selected based on population frequencies in the gnomAD database that would suggest that they were Likely Benign as compared to the incidence of Cohen-Gibson syndrome (COGIS). EED variants deemed as causative for Cohen-Gibson syndrome (P and LP categories) were selected from clinical reports in the literature.

These variants were then prioritised by the following criteria: variant seen in multiple independent individuals, multiple variants reported at the same amino acid position, and/or reported functional data indicating detrimental effects on protein function. EED variants reported in haematological malignancies were chosen if published data had demonstrated an impairment in EED function. An additional EED null variant was also selected, which has been shown to be null in *in vivo* mouse data. Four additional VoUS were tested, as were two component variants derived from the *306R_307NdelinsTD* mutation. See Table 1 for a list of all variants assayed.

### Generation of the knock-in *EED H258Y* (*esc H238Y*) allele

Supporting Figure 2 illustrates the generation of these knock-in fly lines. 3-5 kb of the *esc* gene regions were PCR amplified from the (*y[1] M{w[+mC]=nos-Cas9.P}ZH-2A w[*]*, BL: 5459) fly line and subcloned into pJET2.1 (Thermofisher). gRNAs were designed using the online platform, flyCRISPR target finder (https://flycrispr.org/)^49^ and oligonucleotides with target sequences were inserted into the sgRNA expression vector pUS-BbsI-chiRNA.

esc gRNA 1: CTTCGTTGAAGTCGCGGCCTAATT

esc gRNA 2: /AAACGTGCTGTTCCAAATTATTTC

The homology-directed repair dsDNA template contained the knock-in mutations, PAM mutations, homology arms and 3xP3EGFP transposon cassette. The *esc* mutations and PAM mutations (to prevent re-excision of the modified genomic region) were engineered using primers with the desired mutation. 1kb homology arms were amplified from the pJET vector, and a 3XP3EGFP transposon cassette was amplified by PCR from pScarlessHD-EGFP. Homology arms, DNA template with desired mutations and 3XP3EGFP cassette were assembled by HiFi DNA assembly (NEB). Wildtype esc donor constructs that also included the PAM site mutations were also generated for use as controls. Plasmids were sequence verified and purified with midi prep (Qiagen). Plasmids with gRNA vectors were sent for injection (completed by Genome Prolab Inc. (Quebec).

*esc-Ref* and *H258Y* crispant lines were outcrossed for 6 generations with *w*^1118^ to remove any off-target mutations, and then sequenced to verify presence of PAM site and knock-in mutations.

Independent *esc-Ref* and *H258Y* crispant males were crossed to *w-; ES/CyO* balancer line to generate stable lines of *w-; esc-Ref orH258Y /CyO*. The *3XP3-EGFP* transposon cassette in these lines was located in the 3’UTR and was not predicted to disrupt gene expression. However, the *3XP3-EGFP* cassette was removed from *esc-Ref* and *H258Y.* For *3XP3-EGFP* cassette removal, *w^-^;esc-Ref or H258Y /CyO* males/females were crossed to PiggyBac transposase (males and females). F1 generation GFP mosaic females (*w-;CyO,PBac/esc^#^-3XP3-EGFP;TM6, Tb/+)* were selected and crossed to *w-;ES/CyO* males. F2 generation w*-;esc^#^/CyO* males were crossed to *w-;ES/CyO* females and F3 generation *w-;esc^#^/CyO* males and females were inter-crossed to generate stable lines.

### Construction and Generation of *esc* mimetic Lines by Recombinase Mediated Cassette Exchange (RMCE)

The *esc* cDNA, SV40 polyA and 3XP3dsRed were assembled by HiFi assembly and cloned into the AgeI/HindII site of *pBS-KS-attB2-SA(0)-T2A-GAL4* (DGRC#1409). *esc* variants (*esc* mimetics) were generated with splicing by overlap-extension PCR using primers with desired mutations and cloned into the AgeI or ApaI and EcoR1 site of *pattB-SA(0)-T2A-esc-3XP3dsRed-attB*. Plasmids were injected into *esc::T2A::GAL4* CRIMIC line (BL79282) and the new cassette integrated by ϕC31-integrase mediated transgenesis by Genome Prolab Inc. (Quebec).

Fly lines were returned as stable lines with the *esc* RMCE cassette over a *CyO* balancer chromosome, or as parental males crossed to *w*^1118^*; Adv/CyO* balancer line, from which we generated a stable line over a *CyO* balancer chromosome. All fly lines used in subsequent experiments were sequence-verified. 22 RMCE mimetic lines were generated using this method (Figure 1), and experiments were done in three batches. For convenience in interpretation of the relevance of the fly data to human variants, RMCE mimetic mutations are referred to by the human variant nomenclature.

### Imaging Adult Wings and Legs

Adult flies were collected and placed in isopropanol and stored at 4°C until dissection. Wings and legs were removed and mounted with Canada Balsam on glass slides and imaged using ZEISS SteREO Discovery.V8 with a Huawei rear camera 16 MP, f/2.2, 1/3.1”, 1.0µm, PDAF 2 MP, (depth) or Nikon Z50 camera attached on a Leica MZ16F stereomicroscope. For notum and abdomen pictures, CO2-anaesthetized flies were imaged with a Nikon Z50 camera coupled to a Leica MZ16F stereomicroscope. 4-5 Z-stack images were taken aligned and stack using Zerene Stacker software.

### Statistical Analyses

Expected values for knock-in mutation experimental groups were calculated based on expected Mendelian ratios. For *esc* mimetic and UASz-esc mimetic experimental groups, expected values were calculated based on observed values of the *esc-Ref*. Chi Squared testing and statistical significance values were generated using the online chi-squared calculator by GraphPad.^50^

## Author Contributions

Variant Identification within databases and published literature was done by Dr. Cyrus and Dr. Gibson. Dr. Medina Giró designed the GAL4/UAS rescue based experiment. All other experiments were designed by Drs. Cyrus, Allan, Medina Giró and Gibson, and conducted by Drs. Cyrus and Medina Giró along with analysis and interpretation of results with oversight from Dr. Allan and Dr. Gibson. Molecular work was performed by Drs. Lian and Medina Giró. The manuscript was written and edited by Dr. Cyrus with comments and recommendations by Drs.

Gibson, Allan and Medina Giró. All authors approved the manuscript prior to submission.

## Acknowledgements

Stocks obtained from the Bloomington *Drosophila* Stock Center (NIH P40OD018537) were used in this study. We thank *Drosophila* Genomics Resource Center, supported by NIH grant 2P40OD010949 for the esc cDNA. We used FlyBase (releases 2020 through 2024) to find information on phenotypes/function/stocks/gene expression (etc). We would like to thank Gino Laberge at Genome ProLab for the kind gift of balancer stocks and also their transgenic services (Genome ProLab Inc, Quebec, Canada). Funding for this work was provided by Canadian Institutes of Health Research AWD-013471 (PJT-168982, including CRCEF funding) and Simons Foundation Autism Research Initiative - 2021 Genomics of ASD: Pathways to Genetic Therapies award ID 886158. W.T.G.’s research salary is supported by the BC Children’s Hospital Research Institute through an intramural IGAP award.

