## Supplemental Information for "Mimicking Human EED Variants in Drosophila: A Promising Strategy to Analyse Human EED Variant Function"

### Supplementary Information

Sharri S. Cyrus[a,b], Sònia Medina Giró[c], Tianshun Lian[c], Douglas W. Allan[c,d*], William T. Gibson[a,b*]

a Department of Medical Genetics, University of British Columbia, Vancouver, British Columbia, Canada

b British Columbia Children's Hospital Research Institute, Vancouver, British Columbia, Canada

c Department of Cellular and Physiological Sciences, Life Sciences Institute, University of British Columbia, Vancouver, British Columbia, Canada

d Djavad Mowafaghian Centre for Brain Health, University of British Columbia, Vancouver, British Columbia, Canada

* Douglas W. Allan is the corresponding author regarding *Drosophila* experiments and strains and William T. Gibson is the clinical corresponding author.

| | HOMOZYGOSITY AT ROOM TEMPERATURE | | | HOMOZYGOSITY AT 25°C | | |
|---|---|---|---|---|---|---|
| **Variant** | Expected (%) | Observed (%) | Statistical Significance | Expected (%) | Observed (%) | Statistical Significance |
| **Ref** | - | 66 (33%) | | - | 44 (36%) | |
| **T50P**[LB] | 36 (33%) | 31 (28%) | ns | 46 (36%) | 48 (38%) | ns |
| **I274V**[LB] | 28 (33%) | 32 (28%) | ns | 60 (36%) | 58 (35%) | ns |
| **N194S**[C] | 39 (33%) | 23 (20%) | *** | 35 (36%) | 2 (2%) | **** |
| **R236G**[C] | 38 (33%) | 0 (0%) | **** | 68 (36%) | 0 (0%) | **** |
| **H258Y**[C] | 37 (33%) | 23 (20%) | ** | 54 (36%) | 11 (7%) | **** |
| **R302S**[C] | 31 (33%) | 0 (0%) | **** | 43 (36%) | 0 (0%) | **** |

**Supporting Table 1: Observed and Expected Frequencies of Homozygosity in Flies Bearing Likely Benign and COGIS Mimetic Variants**

Flies heterozygous for the indicated *T2A::esc*[#] mimetic variants were intercrossed and the number of homozygous variant progeny were counted. Data are expressed as number of surviving adults observed, compared to the numbers expected according to Mendelian ratios. Expected (%) are estimated based on the observed *esc::Ref* control values included in each batch. Additionally, on average 33% of flies were expected in the *esc::Ref* group, as parental flies were heterozygous for a balancer allele, that is homozygous adult lethal. Chi-Square testing indicated significant lethality among flies homozygous for COGIS variants, but not among homozygotes for Likely Benign variants.

LB: Likely Benign variant; C: COGIS variant; Chi-Square testing indicates that the difference in expected vs observed frequencies is: significant * $p \geq 0.05$; ** p= 0.001 to 0.01; *** p= 0.0001 to 0.001; **** $p < 0.0001$ or not significant (ns).

| | HOMOZYGOSITY AT ROOM TEMPERATURE | | | HOMOZYGOSITY AT 25°C | | |
|---|---|---|---|---|---|---|
| **Variant** | Females | Males | Totals | Females | Males | Totals |
| **esc-Ref** | 33 | 33 | 66 | 17 | 27 | 44 |
| **T50P[LB]** | 14 | 17 | 31 | 27 | 21 | 48 |
| **I274V[LB]** | 13 | 19 | 32 | 28 | 30 | 58 |
| **N194S[C]** | 11 | 12 | 23 | 1 | 1 | 2 |
| **R236G[C]** | 0 | 0 | 0 | 0 | 0 | 0 |
| **H258Y[C]** | 7 | 16 | 23 | 5 | 6 | 11 |
| **R302S[C]** | 0 | 0 | 0 | 0 | 0 | 0 |

**Supporting Table 2: Total Number of Homozygotes bearing Likely Benign and COGIS-Mimetic Variants (Calibration Phase)**
Heterozygous *T2A::esc*# mimetic variants were intercrossed and the number of homozygous progeny bearing the indicated variant were counted.
LB: Likely Benign variant; C: COGIS variant

| Variant | Expected (%) | Observed (%) | Statistical Significance |
|---|---|---|---|
| **Likely Benign Variants** | | | |
| T50P$^{LB}$ | 225(39%) | 230 (40%) | ns |
| I274V$^{LB}$ | 226 (39%) | 196 (34%) | ** |
| **Likely Pathogenic/ Pathogenic COGIS Variants** | | | |
| N194S$^{C}$ | 210 (39%) | 152 (28%) | **** |
| R236G$^{C}$ | 197 (39%) | 155 (31%) | **** |
| R236T$^{C}$ | 214 (33%) | 218 (34%) | ns |
| D237G$^{C}$ | 224 (33%) | 232 (34%) | ns |
| H258Y$^{C}$ | 217 (39%) | 188 (34%) | ** |
| R302G$^{C}$ | 232 (40%) | 147 (25%) | **** |
| R302S$^{C}$ | 193 (39%) | 149 (30%) | **** |
| 306R_307NdelinsTD$^{C}$ | 227 (33%) | 220 (32%) | ns |
| A378V$^{C}$ | 100 (40%) | 34 (14%) | **** |
| **LoF Variants in Haematological Malignancies** | | | |
| L240Q$^{M}$ | 281 (40%) | 164 (23%) | **** |
| G255D$^{M}$ | 301 (40%) | 92 (22%) | **** |
| S259F$^{M}$ | 258 (40%) | 203 (31%) | **** |
| I363M$^{M}$ | 241 (40%) | 201 (33%) | *** |
| **Variants of Uncertain Significance** | | | |
| R52C$^{V}$ | 296 (40%) | 264 (36%) | * |
| T158M$^{V}$ | 284 (33%) | 254 (36%) | * |
| F198L$^{V}$ | 285 (33%) | 266 (37%) | ns |
| H258L$^{V}$ | 210 (33%) | 156 (24%) | **** |
| **306R_307NdelinsTD Component Variants** | | | |
| R306T$^{V}$ | 224 (39%) | 201 (35%) | * |
| N307D$^{V}$ | 212 (39%) | 207 (38%) | ns |
| N307D$^{V}$ | 95 (33%) | 101 (35%) | ns |
| **Null Variant** | | | |
| L196P$^{N}$ | 228 (40%) | 1 (0.2%) | **** |
| **esc-Ref** | | | |
| esc-Ref (Batch1) | - | 200 (39%) | - |
| esc-Ref (Batch 2) | - | 84 (33%) | - |
| esc-Ref (Batch 3) | - | 165 (40%) | - |

**Supporting Table 3: Observed and Expected Frequencies of Adult Flies Hemizygous for *esc* Mimetic Variants**

Males bearing the noted *T2A::esc*[#] mimetic variants in the heterozygous state were crossed to virgin females heterozygous for *esc-Df* (i.e. virgin females hemizygous for wild type *esc*), and the number of adult progeny hemizygous for each variant were counted. Data are expressed as number (and percentage) of adults observed versus number expected based on Mendelian ratios. LB: Likely Benign variant; C: COGIS variant; M: Haematological malignancy variant; N: null variant; V: Variant of uncertain significance;
Expected (%) are estimated based on the observed esc-Ref control values included in each batch. Data from Table 2 is included in this table for the Likely Benign and COGIS variant subtypes. Chi-Square testing found that the difference in expected vs observed frequencies was: significant * $p \geq 0.05$; ** p= 0.001 to 0.01; *** p= 0.0001 to 0.001; **** $p < 0.0001$ or not significant (ns).

| Variant | Females | Males | Totals |
|---|---|---|---|
| **Likely Benign Variants** | | | |
| T50P[LB] | 124 | 106 | 230 |
| I274V[LB] | 108 | 88 | 196 |
| **Likely Pathogenic/ Pathogenic COGIS Variants** | | | |
| N194S[C] | 75 | 77 | 152 |
| R236G[C] | 68 | 87 | 155 |
| R236T[C] | 108 | 110 | 218 |
| D237G[C] | 117 | 115 | 232 |
| H258Y[C] | 90 | 98 | 188 |
| R302G[C] | 68 | 79 | 147 |
| R302S[C] | 77 | 72 | 149 |
| 306R_307NdelinsTD[C] | 96 | 124 | 220 |
| A378V[C] | 18 | 16 | 34 |
| **LoF Variants in Haematological Malignancies** | | | |
| L240Q[M] | 60 | 104 | 164 |
| G255D[M] | 15 | 77 | 92 |
| S259F[M] | 101 | 102 | 203 |
| I363M[M] | 103 | 98 | 201 |
| **Variants of Uncertain Significance** | | | |
| R52C[V] | 130 | 134 | 264 |
| T158M[V] | 123 | 131 | 254 |
| F198L[V] | 129 | 137 | 266 |
| H258L[V] | 75 | 81 | 156 |
| **306R_307NdelinsTD Component Variants** | | | |
| R306T[V] | 101 | 100 | 201 |
| N307D[V] | 101 | 106 | 207 |
| N307D[V] | 50 | 51 | 101 |
| **Null Variant** | | | |
| L196P[N] | 0 | 1 | 1 |
| **esc-Ref** | | | |
| esc-Ref (Batch1) | 102 | 98 | 200 |
| esc-Ref (Batch 2) | 48 | 36 | 84 |
| esc-Ref (Batch 3) | 77 | 88 | 165 |

**Supporting Table 4: Total Number of Viable Hemizygotes Bearing *esc* Mimetic Variants**
Males heterozygous for each of the above *T2A::esc*[#] mimetic variants were crossed to virgin females heterozygous for *esc-Df*, and the number of adult progeny that were hemizygous for each mimetic variant were counted.

| Variant | Developmental Stage Reached | Adult Progeny with Wing and Notum Abnormalities | Total Adult Progeny |
|---|---|---|---|
| **Likely Benign Variants** | | | |
| T50P[LB] | Adult | 0 | 206 |
| I274V[LB] | Adult | 0 | 168 |
| **Likely Pathogenic/ Pathogenic COGIS Variants** | | | |
| N194S[C] | Embryonic/Early Larval Lethal | N/A | N/A |
| R236G[C] | Pupal/Late Larval Lethal | N/A | N/A |
| R236T[C] | Pupal/Late Larval Lethal | N/A | N/A |
| D237G[C] | Adult | 23 | 352 |
| H258Y[C] | Adult | 15 | 23 |
| R302G[C] | Embryonic/Early Larval Lethal | N/A | N/A |
| R302S[C] | Embryonic/Early Larval Lethal | N/A | N/A |
| 306R_307NdelinsTD[C] | Adult | 13 | 34 |
| A378V[C] | Adult | 9 | 40 |
| **LoF Variants in Haematological Malignancies** | | | |
| L240Q[M] | Adult | 35 | 88 |
| G255D[M] | Embryonic/Early Larval Lethal | N/A | N/A |
| S259F[M] | Adult | 37 | 198 |
| I363M[M] | Adult | 6 | 151 |
| **Variants of Uncertain Significance** | | | |
| R52C[V] | Adult | 0 | 365 |
| T158M[V] | Adult | 0 | 435 |
| F198L[V] | Adult | 0 | 528 |
| H258L[V] | Embryonic/Early Larval Lethal | N/A | N/A |
| **306R_307NdelinsTD Component Variants** | | | |
| R306T[V] | Adult | 0 | 219 |
| N307D[V] | Adult | 22 | 97 |
| N307D[V] | Adult | 39 | 242 |
| **Null Variant** | | | |
| L196P[N] | N/A | N/A | N/A |
| **esc-Ref** | | | |
| esc-Ref (Batch1) | Adult | 0 | 171 |
| esc-Ref (Batch 2) | Adult | 0 | 141 |
| esc-Ref (Batch 3) | Adult | 0 | 116 |

**Supporting Table 5: Maternal Effects On Survival Observed Among Progeny of Females Hemizygous for *esc* Mimetic Variants**

Adult female virgins hemizygous for each indicated variant were crossed to $w^{1118}$ males, and the developmental stage reached by the progeny was recorded. Adult progeny with visibly abnormal phenotypes were included with apparently normal adult progeny in the "Total Adult Progeny" column. Information in this table is included in Figure 6B.

| | | *esc*$^{\#}$/ *esc::T2A::GAL4* | | *esc*$^{\#}$/*esc-Df* | | *esc*$^{\#}$/*esc*$^{5}$ | |
|---|---|---|---|---|---|---|---|
| | **LINE** | Expected | Observed | Expected | Observed | Expected | Observed |
| ***esc-Ref*$^{KI}$** | **1** | 119 | 111 | 164 | 190 | 11 | 7 |
| | **2** | - | - | 190 | 215 | 12 | 19 |
| | **3** | - | - | - | - | 17 | 15 |
| | **4** | 75 | 77 | - | - | - | - |
| | **5** | 62 | 56 | 163 | 188 | 11 | 11 |
| | | 255 (50%)[a] | 244 (48%) | 517 (33%) | 593 (38%) | 51 (33%) | 52 (34%) |
| ***esc-H258Y*$^{KI}$** | **1** | 43 | 55 | 149 | 118 | 16 | 10 |
| | **2** | - | - | 149 | 44 | 11 | 2 |
| | **3** | 64 | 60 | - | - | - | - |
| | **4** | 69 | 63 | 123 | 50 | 10 | 3 |
| | | 176 (33%) | 178 (33%) [ns] | 421 (38%) | 212 (19%) [****] | 37 (33%) | 15 (13%) [****] |

**Supporting Table 6 Observed and Expected Frequencies of *H258Y* Compound Heterozygote and Hemizygote Crispants**

At 25°C, heterozygous *esc-H258Y* males or homozygous/heterozygous *esc-Ref males* were crossed to virgin females heterozygous for one of three alleles: *esc::T2A::GAL4*, *esc-Df* or *esc*$^{5}$. The table shows the expected Mendelian ratios, as well as the observed ratios for *esc-Ref* and *esc-H258Y* compound heterozygote and hemizygote progeny. '*' Chi-Square testing indicates significant lethality prior to adulthood for *esc-H258Y/esc-Df and for esc-H258Y/esc*$^{5}$. Chi-Square testing indicates that the difference in expected vs observed frequencies is: significant * $p \geq 0.05$; ** p= 0.001 to 0.01; *** p= 0.0001 to 0.001; **** $p < 0.0001$ or not significant (ns)

| | | $esc^{\#}$/ esc::T2A::GAL4 | | $esc^{\#}$/esc-Df | | $esc^{\#}/esc^{5}$ | |
|---|---|---|---|---|---|---|---|
| | **LINE** | Females | Males | Females | Males | Females | Males |
| ***esc-Ref*** | **1** | 69 | 42 | 105 | 85 | 4 | 3 |
| | **2** | - | - | 104 | 111 | 13 | 6 |
| | **3** | - | - | | | 7 | 4 |
| | **4** | 35 | 42 | | | - | - |
| | **5** | 33 | 23 | 98 | 90 | 9 | 6 |
| | | 137 | 107 | 307 | 286 | 33 | 19 |
| ***esc-H258Y*** | **1** | 21 | 34 | 70 | 48 | 9 | 1 |
| | **2** | - | - | 27 | 17 | 1 | 1 |
| | **3** | 36 | 24 | | | - | - |
| | **4** | 37 | 26 | 36 | 14 | 1 | 2 |
| | | 94 | 84 | 133 | 79 | 11 | 4 |

**Supporting Table 7: Total number of females and males per genotype of *H258Y* Compound Heterozygote and Hemizygote Crispants**

At 25°C heterozygous *esc-H258Y* males or homozygous/heterozygous *esc-Ref males* were crossed to virgin females heterozygous for one of three alleles: *esc::T2A::GAL4*, *esc-Df* or $esc^{5}$. The table shows the total number of adult flies of the above genotypes recovered from the cross.

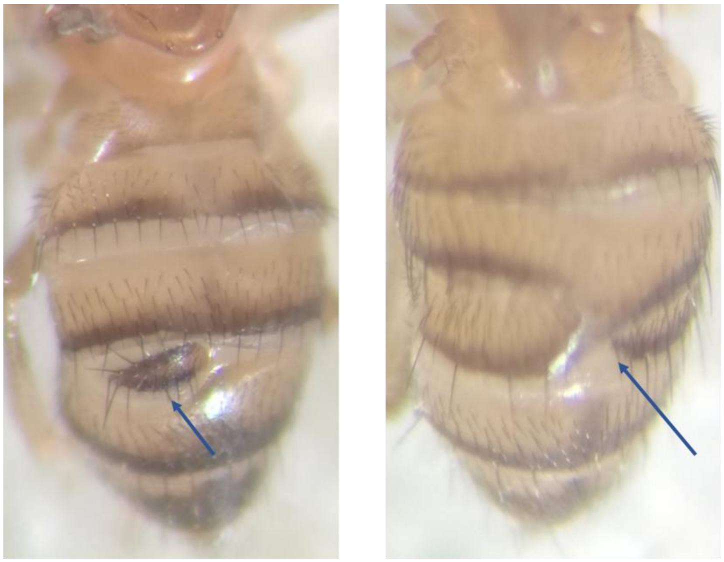

**Progeny of *esc-Ref/esc::T2A::GAL4*** **Progeny of *T50P/esc::T2A::GAL4***

Supporting Figure 1: Representative Images of Tergite Defects Observed in Flies

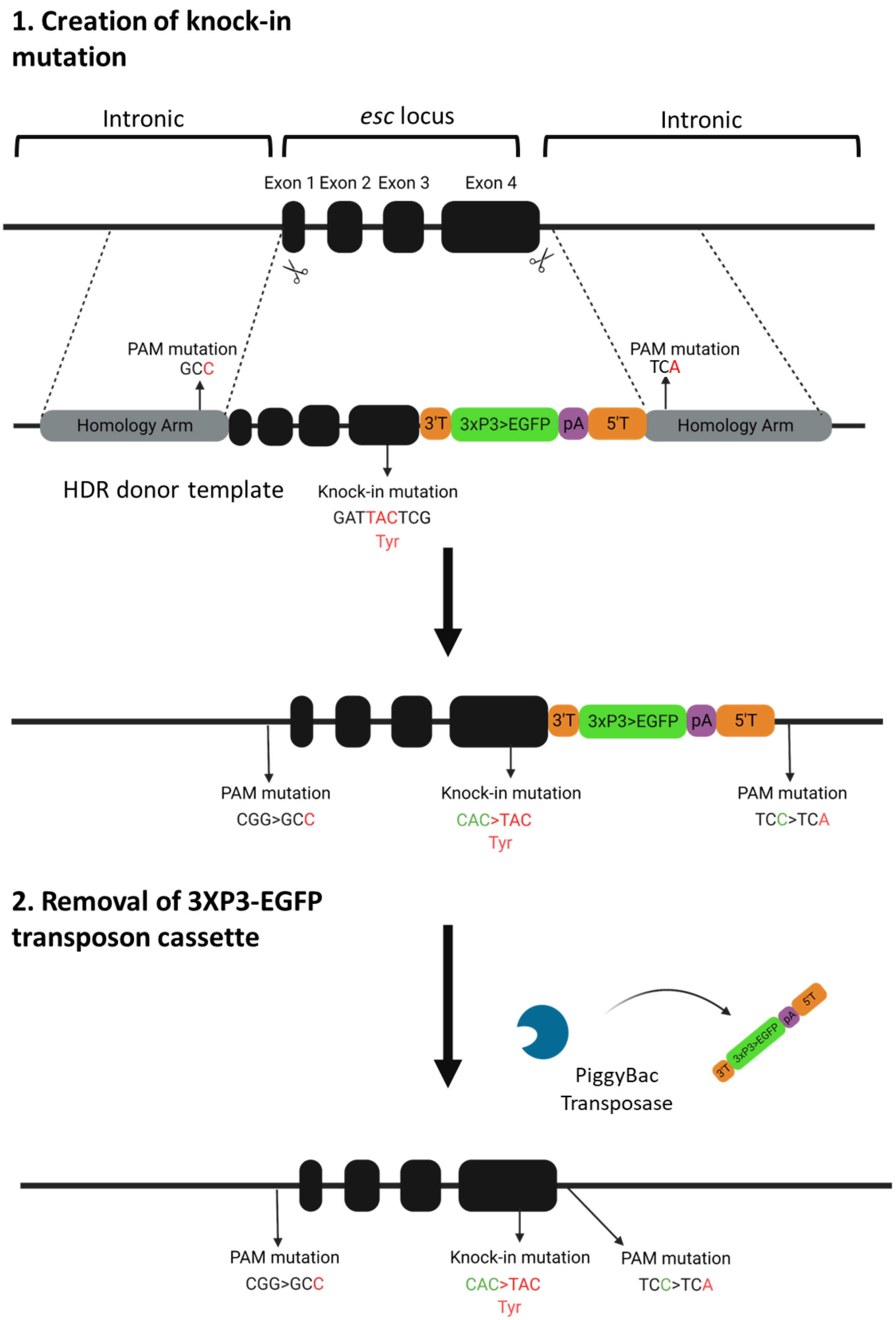
1. Creation of knock-in mutation
Intronic
esc locus
Intronic
Exon 1
Exon 2
Exon 3
Exon 4
PAM mutation
GCC
PAM mutation
TCA
Homology Arm
3'T
3xP3>EGFP
pA
5'T
Homology Arm
HDR donor template
Knock-in mutation
GATTACTCG
Tyr
3'T
3xP3>EGFP
pA
5'T
PAM mutation
CGG>GCC
Knock-in mutation
CAC>TAC
Tyr
PAM mutation
TCC>TCA
2. Removal of 3XP3-EGFP transposon cassette
PiggyBac
Transposase
3'T
3xP3-EGFP
pA
5'T
PAM mutation
CGG>GCC
Knock-in mutation
CAC>TAC
Tyr
PAM mutation
TCC>TCA

**Supporting Figure 2 Generation of knock-in EED H258Y (esc H238Y) mutation by CRISPR/Cas9**
**Creation of knock-in mutation:** Guide RNAs directed Cas9 to genomic target, where it created double strand breaks at genomic sites indicated by the scissors. Homology directed repair (HDR) occured using the HDR donor template. The HDR donor template contained the homology arms with protospacer-adjacent motif (PAM) mutations, the entire *esc* locus sequence with esc H238Y mutation and transposon cassette: 3'Transposon; 3xP3-EGFP: green eye fluorescent marker; pA: SV40pA terminator sequence. HDR results in the creation of the knock-in esc H238Y mutation and insertion of the transposon cassette (3'T-3xP3-EGFP-pA-5'T). PAM mutations stopped subsequent cleavage by Cas9, thereby preventing excision of the newly modified genomic region. They occur in an intronic region of the *Pde1c* gene and are not predicted to disrupt gene expression.
**Removal of 3xP3-EGFP transposon cassette:** The PiggyBac transposase line efficiently excised the transposon cassette. Image Created with BioRender.com

#### Supplemental Methods

##### Estimation of the Collective Prevalence of Rare Coding Variation in PRC2 Core Complex members EED, EZH2 and SUZ12 among gnomAD Participants

We sought to estimate the prevalence of rare coding variants among the general population. Since all human genetic databases are subject to some level of recruitment bias, there is no extant database that represents the "general population." Nevertheless, the gnomAD database version 4.0 release contains 730,947 exomes and 76,215 genomes (total 807,162 individuals mapped to the GRCh38/hg38 human genome build), from a variety of ancestral backgrounds.

The **gnomAD browser** (https://gnomad.broadinstitute.org/) was consulted November 16, 2023 and one of us (W.T.G.) downloaded gnomAD variants meeting classification criteria "pLoF" and Missense/Inframe indel for the genes EED, EZH2 and SUZ12. Variants under the "Synonymous" and "Other" categories were not investigated, nor were structural variants. gnomAD's "Transcript view" was set to the current Matched Annotation from NCBI and EMBL-EBI (MANE) transcript for each gene (ENST00000322652.10 for EED, ENST00000320356.7 for EZH2 and ENST00000322652.10 for SUZ12).

Variants annotated with the ClinVar Clinical Significance tags as "Benign" or "Likely Benign" were excluded from the calculation of Aggregate Rare Variant MAF; because these are least likely to affect protein function, they are least likely to have health consequences and thus to be of interest for aggregate rare variant burden tests. Typically, variants with minor allele frequencies that are high enough to merit the descriptors "Benign" or "Likely Benign" would be best assessed for disease relevance using Genome-Wide Association Studies (GWAS). Low-frequency variants that are predicted by software to be Likely Benign (such as rs972308238) may or may not have functional effects – exclusion of these variants from estimates of Aggregate Rare Variant MAF will not have much effect on the magnitude of the estimate.

Summation of the gnomAD-derived MAFs shows that, taken together, rare coding variants in the PRC2 "core" complex are expected in approximately 1.3% of chromosomes, or 2.6% of the population. The signal appears largely driven by rare variants in SUZ12. Also worthy of note is the fact that rare PRC2 variants classified as Pathogenic and Likely Pathogenic are observed in gnomAD data, albeit at extremely low frequencies. Pathogenic EED variants p.Asn194Ser and p.Arg302Gly were observed in 2 and 1 participants, respectively. Likely Pathogenic EZH2 variants p.His129Arg, p.Pro577Leu, p.Thr683Asn and p.Arg690His were observed once each, and Pathogenic EZH2 variant p.Val626Met was seen twice. Likely Pathogenic SUZ12 variants p.Leu385ProfsTer10 and Pathogenic SUZ12 variant p.Arg654Ter were also observed once each.

| PRC2 Member | Aggregate MAF<br>Rare missense and in-frame Indel Variants | Aggregate MAF<br>Rare predicted Loss-of-Function Variants | Overall Total Aggregate MAF |
|---|---|---|---|
| **EZH2** | 0.002622 | 0.000074 | 0.002696 |
| **EED** | 0.002083 | 0.000028 | 0.002111 |
| **SUZ12** | 0.008027 | 0.000311 | 0.008338 |
| **PRC2 Core** | 0.012732 | 0.000413 | 0.013145 |

The **Genome Asia Browser** (https://www.genomeasia100k.org/) was consulted October 31, 2023 and one of us (W.T.G.) created worksheets similar to the above by manual inspection of the variant lists obtained by gene- and transcript-specific queries with inclusion criteria "Missense + LoF." Summation of the Genome Asia-derived MAFs as of shows that, taken together, rare coding variants in the PRC2 “core” complex are expected in approximately 1% of chromosomes, or 2% of the Asian population, based on a non-random heterogeneous sample of ~3300 chromosomes from individuals annotated in the database as of 31 October, 2023.

| PRC2 Member | Aggregate MAF Rare missense and in-frame Indel Variants | Aggregate MAF Rare predicted Loss-of-Function Variants | Overall Total Aggregate MAF |
|---|---|---|---|
| **EZH2** | 0.0009 | Not Observed | 0.0009 |
| **EED** | 0.0043 | Not Observed | 0.0043 |
| **SUZ12** | 0.0039 | 0.0006 | 0.0045 |
| **PRC2 Core** | 0.0091 | 0.0006 | 0.0097 |

The **MSSNG Browser** (https://research.mss.ng/) was consulted October 31, 2023. It contains 11,364 human genomes, of which 5,134 (~45%) are derived from persons with Autism Spectrum Disorders (ASD). The DB6 release became available October 16, 2019 and is based on the GRCh38/hg38 human genome build. Participants in this database are highly selected for autism and other neurodevelopmental disorders, with most participants who do not have these disorders consisting of unaffected relatives of participants with these phenotypes. Though variant frequencies in the MSSNG data are not representative of samples of the “general” (i.e. randomly-selected) human population, rare allele frequencies in neurodevelopmentally-important genes may be informative of the genetic and allelic architecture of these neurotypes.

After completion of the appropriate data access agreements, one of us (W.T.G.) queried the MSSNG Browser for rare coding variants in EZH2, EED and SUZ12. The parameters of the query included the relevant gene Symbol, Variant quality (Passing), Affection (affected), Control Database Frequency (<= 0.01) and Damage Potential (High). Query results were exported as .tsv files and converted to .xlsx files in a manner similar to that done for queries of the gnomAD database above.

Within the MSSNG database, the number of individuals carrying heterozygous variants in EZH2, EED and SUZ12 that satisfied the above queries (Benign/Likely Benign variants excluded) was 30 for EZH2, 3 for EED and 121 for SUZ12. Though the crude total of 154 aggregate variants in the PRC2 core complex members is difficult to interpret among the 5,134 MSSNG participants (since some affected sib pairs are included), we may at least conclude that rare coding variants predicted to affect PRC2 complex function to some extent are commonly seen among children and adults with neurodevelopmental disorders that include autism endotypes.
